# The potential of AI models to identify regulatory variants underlying cattle traits

**DOI:** 10.1101/2024.08.01.606140

**Authors:** R Zhao, R Owen, M Marr, S Jayaraman, N Chue Hong, A Talenti, MA Hassan, JGD Prendergast

**Author notes:** Corresponding authors RZ JGDP.

## Abstract

Considerable progress has been made in developing machine learning models for predicting human functional variants, but progress in livestock species has been more limited. This is despite the disproportionate potential benefits such models could have to livestock research, from improving breeding values to prioritising functional variants at trait-associated loci. A key open question is what datasets and modelling approaches are most important to close this performance gap between species. In this work we have developed a new framework for predicting regulatory variants, that includes deriving key conservation metrics in cattle for the first time, variant annotation and model training. When trained on expression quantitative trait loci (eQTL) and massively parallel reporter assay (MPRA) data for human and cattle we show that this framework has a high performance at predicting regulatory variants, with a maximum area under the receiver operating characteristic curve (AUROC) score of 0.86 in human and 0.81 in cattle. We explore various approaches to further close this performance gap, including integrating advanced DNA sequence models and generating extra chromatin data, but illustrate that the best approach would be generating improved gold-standard sets of known cattle regulatory variants. Importantly we demonstrate both the human and cattle models substantially enrich for variants linked to important traits, with up to 18-fold enrichments for functional variants observed. Consequently, with our framework developed to be applicable across species these results not only demonstrate its potential utility for fine-mapping functional variants and improving breeding values in cattle, but also its potential for wider use across animal species.

## Introduction

Regulatory variants play a critical role in shaping mammalian phenotypes and are estimated to account for approximately 60% of the genetic differences underlying human disease (Nica & Dermitzakis, 2013) and around 70% of the heritability for many cattle phenotypes (Xiang et al., 2023). Genome-wide association studies (GWAS) have successfully linked numerous genomic regions to various diseases and traits in both humans and livestock; however, pinpointing the specific causal variants within these regions remains challenging. One major obstacle is linkage disequilibrium (LD) among nearby variants, that is particularly high in livestock breeds, and which makes it difficult to isolate the genetic changes driving the trait. Although intersecting GWAS findings with known expression quantitative trait loci (eQTL) helps suggest regulatory variation as a plausible cause of a GWAS peak, the precise causal variant still often remains elusive.

To overcome this challenge there has been growing interest in developing artificial intelligence (AI) models for predicting functional regulatory variants, though primarily in the human genome. One early example of such an AI model was GWAVA (Ritchie et al., 2014), which employed a random forest model trained on regulatory variants sourced from the human gene mutation database (HGMD) (Stenson et al., 2020). GWAVA leveraged a broad set of genomic features, including chromatin data and sequence conservation, to classify new variants, with an area under the receiver operating characteristic curve (AUROC), a combined measure of the model’s performance across classification thresholds, ranging from 0.75 to 0.84 depending on the chosen background set of non-functional variants. DeepSEA (Zhou & Troyanskaya, 2015) used a convolutional neural network (CNN) to predict chromatin profiles from local sequence and demonstrated that these predictions could be used to classify regulatory variants. Using the predicted effects of sequence changes on chromatin profiles, its models achieved an AUROC of up to 0.7 and outperformed GWAVA when both were trained on the same HGMD data. More recently, DeepSEA’s foundational work was extended to include 21,907 chromatin profiles covering over 1,300 cell lines and tissues, culminating in the Sei model (K. M. Chen et al., 2022). Mutations in HGMD had Sei functional effect scores up to 6.5 times higher than background variants. Another key development is the Enformer model (Avsec et al., 2021), which, similar to DeepSEA, leverages CNNs to predict large-scale omics data from sequence. Training random forest classifiers on Enformer output to distinguish fine-mapped GTEx (GTEx Consortium, 2020) regulatory variants from background variants yielded an AUROC of 0.75.

Despite these advances, most of these efforts to predict functional regulatory variants have focused on humans due to the abundant availability of omics data. Yet extending similar methods to livestock species is of considerable importance. Selective breeding programs in livestock could potentially incorporate functional variant predictions to accelerate genetic gains and yield tangible societal benefits more rapidly than would be feasible for many other species. For example, previous cattle studies found that prioritising variants based on functional-and-evolutionary trait heritability (FAETH) scores (Xiang et al., 2019), encompassing a range of regulatory and evolutionary features, had the potential to enhance the rate of genetic improvement for economically important phenotypes (Xiang et al., 2021). Simulation studies further support the idea that enriching for functional variants can help improve genomic estimated breeding values (Santana et al., 2023). Consequently, identifying functional variants in livestock has the possibility to spur both economic and global public health advances.

A key hurdle to developing advanced machine learning models for livestock species is the scarcity of high-quality training data. In humans, the genotype-tissue expression (GTEx) project has become a gold-standard resource for mapping gene expression and refining genomic regions to fine-mapped regulatory variant using tools such as CaVEMaN (Brown et al., 2017). Inspired by GTEx, the farm animal genotype-tissue expression (FarmGTEx) project aims to establish similarly comprehensive resources in key farm animals, including pigs and cattle (Liu et al., 2022; Teng et al., 2024). For cattle specifically, the FarmGTEx project analysed 7,180 publicly available RNA-Seq samples derived from 46 breeds (or breed combinations) across 114 distinct tissues. eQTL analyses were subsequently carried out for 23 of these tissues. While these collective datasets have significantly advanced eQTL mapping in livestock, challenges remain. For example, the FarmGTEx cattle resource had to integrate heterogenous public datasets generated by different labs at different times, and variant calling was performed solely on RNA-Seq data - ultimately limiting the number of variants detected.

Another major challenge in applying machine learning approaches to livestock species is the relative paucity of specialised functional annotation tools and datasets. Variant annotation is central to genomic analysis, providing functional context to DNA variants. Widely used resources include ANNOVAR (Wang et al., 2010), based on the UCSC Genome Browser database, and the Ensembl Variant Effect Predictor (VEP) (McLaren et al., 2016). Although these tools can deliver a wealth of annotation metadata, they often lack species-specific or downstream-analysis–oriented features, especially those relevant to machine learning approaches for regulatory variant prediction. They can also be human-centric, further underscoring the need for annotation frameworks that are readily adaptable to livestock. Likewise, conservation-based metrics such as PhyloP and PhastCons (Pollard et al., 2010), that have been shown to be invaluable for uncovering functional elements in the human genome (Sullivan et al., 2023), are not currently available for cattle and many other farm animals.

In this study, we sought to assess the feasibility of developing machine learning models that predict regulatory variants in livestock, with a particular focus on cattle. Our first step was to create a species-agnostic variant annotation pipeline tailored for developing machine learning models. This included deriving key PhyloP and PhastCons conservation metrics for cattle for the first time. We then benchmarked prospective modelling approaches by first applying them to human data and comparing that performance to analogous models in cattle. This comparison allowed us to identify potential performance gaps and clarify which new datasets and methods may be most critical for future improvements in cattle regulatory variant prediction. We investigated the effectiveness of different training approaches and feature sets, and the inherent characteristics associated with a variant being predicted as regulatory. Finally, we highlight the substantial utility of both human and cattle models at predicting potential regulatory variants underlying economically and clinically relevant traits.

## Results

### Novel variant annotation pipeline and conservation metrics

To overcome the limitations of existing variant annotation tools for non-human species, we developed *nf-VarAnno*; a species-agnostic and reusable variant annotation pipeline built with Nextflow (Di Tommaso et al., 2017) (Figure 1; https://github.com/evotools/nf-VarAnno). As summarised in Table 1, the pipeline offers five categories of annotations: (1) sequence conservation metrics, (2) variant position properties, (3) VEP annotations, (4) sequence context and (5) predicted functional genomic scores derived from the Enformer deep learning sequence-based model (Avsec et al., 2021).

**Figure 1.**
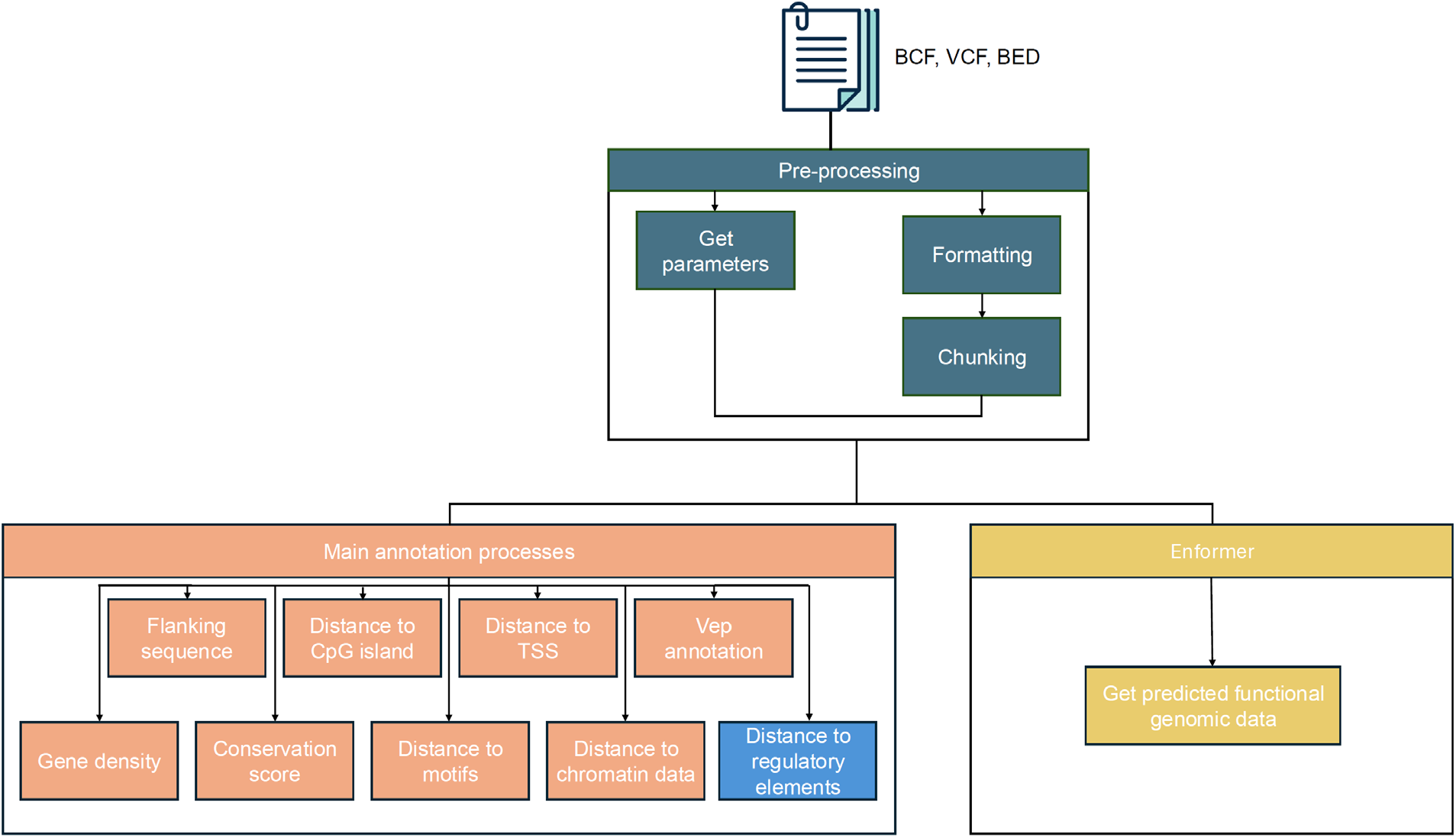
The workflow for the annotation pipeline. The workflow consists of three sections, pre-processing, the main annotation sub-workflow, and Enformer sub-workflow. The blue rectangles indicate annotations specific to humans.

**Table 1.**
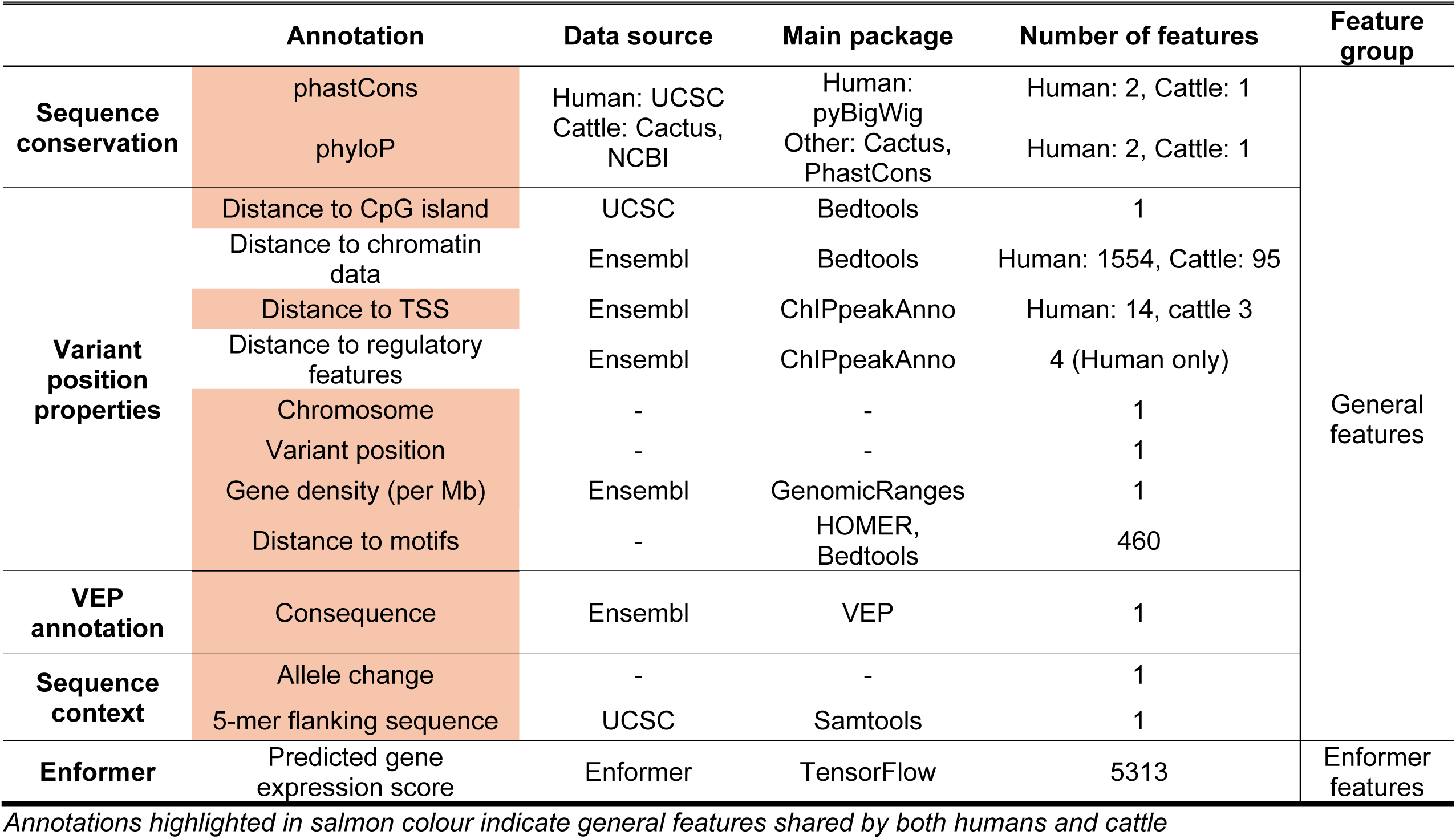
Variant annotation information.

Among these annotation types, sequence conservation metrics are particularly powerful annotations for identifying functional variation in the genome (Sullivan et al., 2023; Xiang et al., 2019). However, aside from GERP (Davydov et al., 2010), most major conservation scores are not available for livestock. To address this gap, we calculated phastCons and phyloP conservation scores for cattle using the Cactus 241-way multiple alignment of vertebrates generated by the Zoonomia consortium (Armstrong et al., 2020; Genereux et al., 2020). As shown in Supplementary Fig. 1A and B, these newly derived metrics are enriched in coding regions, validating their ability to identify functional genomic regions. We have made the full set of cattle conservation metrics publicly available at Zenodo (https://zenodo.org/doi/10.5281/zenodo.13332540).

### The conservation of regulatory variant annotations across species

We next applied our species-agnostic variation annotation pipeline to both human and cattle datasets. In humans, we started with 590,778 regulatory variants identified through CaVEMaN fine-mapping of GTEx data, among which 45,987 exhibited a CaVEMaN causal probability greater than 0.5 and were classified as high-confidence regulatory variants. For cattle, we obtained 79,215 cis-regulatory variants from the cattle FarmGTEx project. To facilitate a direct comparison of annotations between regulatory and background variants, we randomly sampled a background set of variants in each species within 1Mb of transcription start sites (TSS) – consistent with the GTEx cis-eQTL detection window – and then matched their minor allele frequency (MAF) distributions to those of the foreground regulatory variants (See Methods).

Figure 2 demonstrates that many annotation features exhibit similar associations with regulatory variants across species. For example, regulatory variants in both humans and cattle are more likely to involve a G to A (C to T) allele change (Figure 2A, B), and the flanking sequences have a tendency to harbor C:G base pairs (Figure 2I,J). Furthermore, regulatory variants are more abundant in gene-dense regions and exhibit similar enrichments in proximity to specific genomic elements, such as CpG islands and transcription start sites (TSS) (Figure 2C to H). Notably, they also tend to occupy more evolutionarily conserved genomic sites, on average, than the background variants (Supplementary Fig. 2). Taken together, these observations highlight key commonalities in the genomic and evolutionary features of regulatory variants across different species.

**Figure 2.**
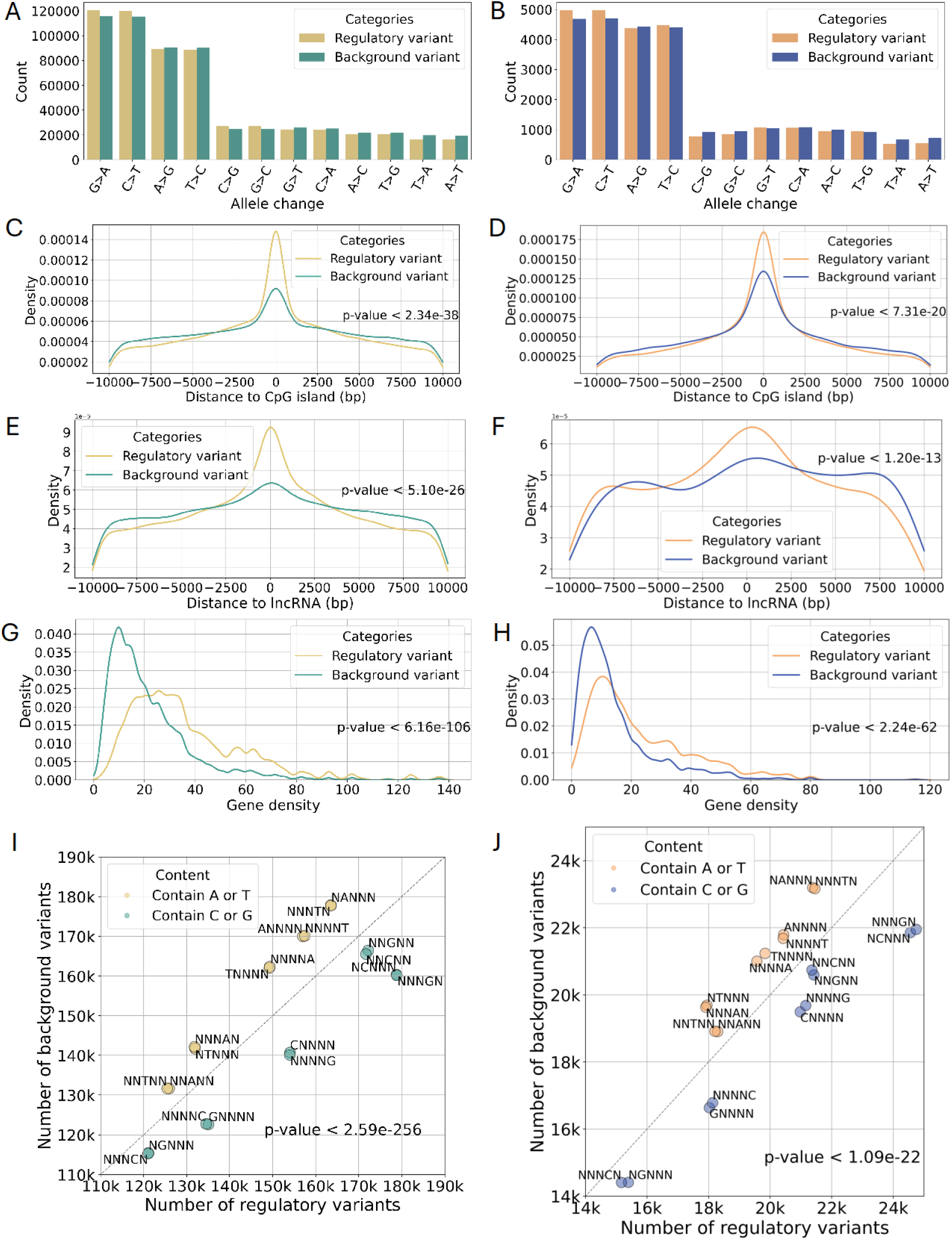
Common characteristics of human and cattle regulatory variants. Number of regulatory and background variants by their allele changes in (A) human and (B) cattle. Kernel density estimate (KDE) plots of distances from regulatory and background variants to CpG island within 10,000bp in (C) human and (D) cattle. KDE plots of distances from regulatory and background variants to lncRNA within 10,000bp in (E) human and (F) cattle. KDE plots of local gene density (genes/Mb) around regulatory and background variants in (G) human and (H) cattle. These annotations display significant differences between the groups in both human and cattle (Two-sample Kolmogorov-Smirnov test p-values are shown in the corresponding plots). Number of variants with different 5-mer flanking sequences between the regulatory and background data in (I) human and (J) cattle. Each circle represents a type of 5-mer flanking sequence with a specific base at a particular position, and the colour of the circle indicates the sequence context. All types of 5-mer flanking sequences exhibit significant differences between the groups in both human and cattle (Chi-Squared test, p-values are shown in the plots).

### The ability to predict regulatory variants in humans

To determine a baseline for using this annotation workflow and set of features to predict regulatory variants, we first applied it to a set of high confidence human regulatory variants. This human model effectively provides a glimpse into the potential future performance of cattle models if trained on the same datasets. The high-confidence human regulatory SNPs were those from the human GTEx project with a CaVEMaN causal probability exceeding 0.5 (45,987 variants) in any tissue. A matching set of background variants was selected from the remaining variants tested in GTEx (See methods). The different sets of variants used in this study can be found in Supplementary Fig. 3. These variants were annotated with general features as indicated in Table 1, excluding the Enformer features that are discussed in a later section. The models were evaluated using a chromosome-level holdout strategy. Specifically, certain chromosomes were set aside as test sets to assess model performance while the remaining chromosomes were used for hyper-parameter tuning (See Methods). Here, we present the model performance tested on chromosomes 1 and 22 (chr1 and chr22). Other test results can be found in Supplementary Fig. 4. The key feature distributions between foreground and background data in the training and test sets are shown in Supplementary Fig. 5. As illustrated in Supplementary Fig. 5, the training and test sets exhibit similar feature distributions for foreground and background data. Different machine learning strategies, including Random Forest, CatBoost, XGBoost, and SVM, were explored, achieving test AUROC scores of 0.82, 0.86, 0.83 and 0.64, respectively, at distinguishing fine-mapped regulatory variants from matching background variants. Considering both its low training time and higher accuracy, we consequently selected the CatBoost model as the algorithm for all downstream analyses.

Leveraging the TraitGym (Benegas, Eraslan, et al., 2025) curated set of putative functional regulatory variants linked to human complex traits, we further benchmarked how well this modelling approach and set of features distinguish human regulatory changes from background variants. The resulting performance (average test AUROC of 0.67 with a standard deviation of 0.04) was broadly comparable to that of the latest version of CADD (∼0.69) (Schubach et al., 2024) the best performing model on this dataset in a recent benchmark (Benegas, Eraslan, et al., 2025), despite CADD’s reliance on a far more extensive set of features - many of which that would be hard to obtain for training livestock models. Performance was also comparable to the recently released GPN-MSA DNA language model (AUROC of ∼0.66) (Benegas, Albors, et al., 2025). Notably, these results were obtained even though one of the potentially most informative biological attributes for classifying regulatory variants, distance to promoters, is effectively controlled for in the TraitGym dataset with the foreground and background variant sets matched on this feature.

Considering that the impact of regulatory variants is typically tissue-specific, we also conducted model training and testing across individual tissues. For this, fine-mapped human variants from five tissues relevant to cattle research (blood, liver, adipose, muscle, and breast) were retrieved from the human CaVEMaN dataset. Due to potential shortages in training data for these tissue-specific datasets, all CaVEMaN fine-mapped variants were used, irrespective of causal probability. Subsequently, the matching background variants were down-sampled in each tissue. As a result, the adipose, liver, blood, breast, and muscle sets comprised 64,378, 24,416, 48,774, 42,022, and 54,244 variants, respectively (each set includes both regulatory and background variants). Among all models, the high-confidence model achieved the highest test AUROC of 0.86 (Figure 3A), and the highest mean cross-validation AUROC score of 0.87 (Supplementary Fig. 4A). To assess the robustness of the high-confidence model, we repeated training and testing 10 times using different randomly selected matching background sets. The models yielded test AUROCs ranging from 0.85 to 0.87, with a mean test AUROC of 0.86 and a standard deviation of 0.006. Although direct comparisons to the performance of other models trained on different datasets should be done with caution, these results compare favourably to other published human models (Avsec et al., 2021; Ritchie et al., 2014; Zhou & Troyanskaya, 2015) and highlights the utility of our workflow for training models that substantially enrich for regulatory variants.

**Figure 3.**
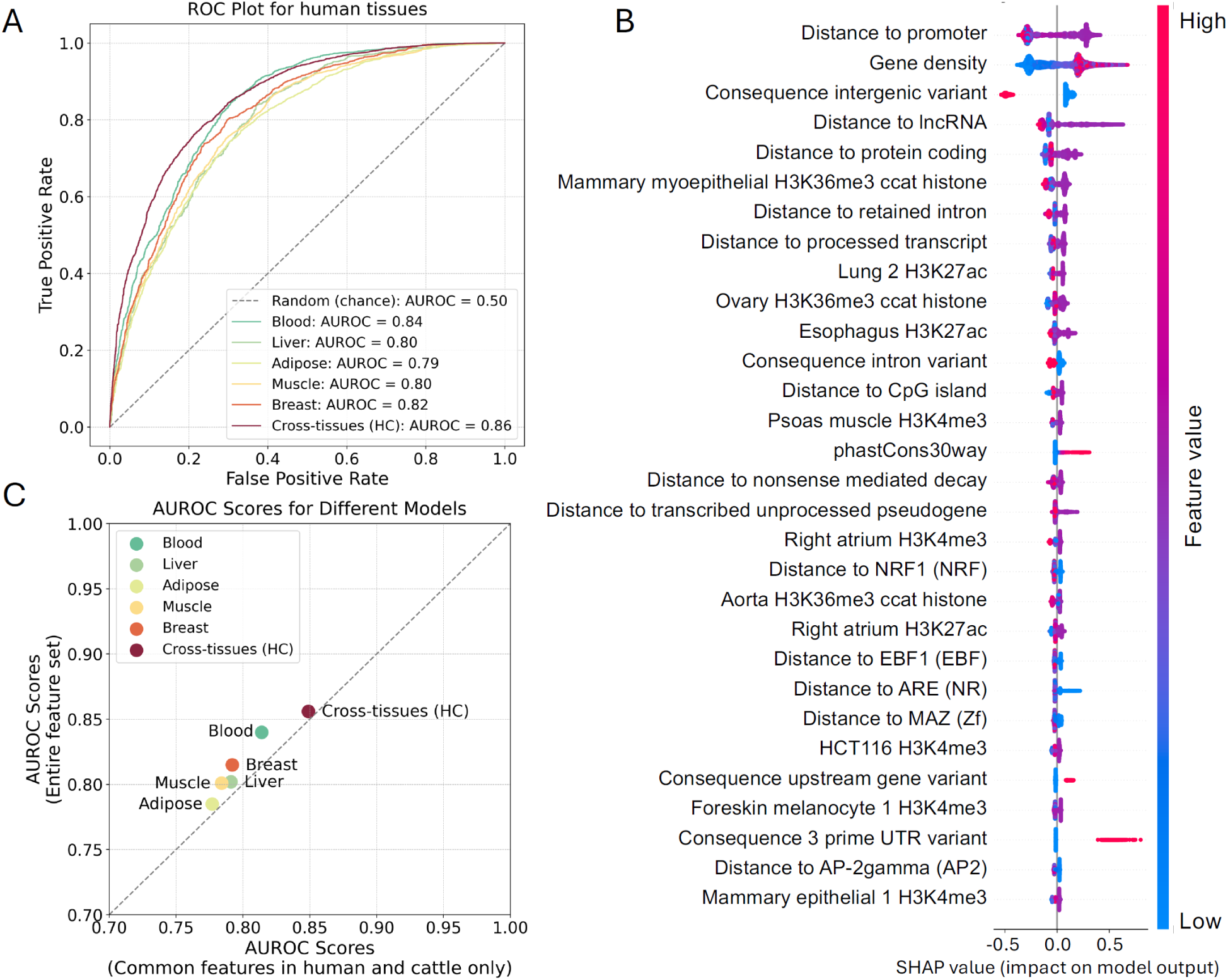
Human model performances. (A) Receiver Operating Characteristic (ROC) curves for human tissue-specific and cross-tissues high-confidence (HC) model. The numbers in the plot represent the area under the receiver operating characteristic (AUROC) scores for different models (ranging from 0 to 1), which evaluate the models’ ability to distinguish regulatory variants. An AUROC closer to 1 indicates a better model. (B) The SHapley Additive exPlanations (SHAP) plot for the human high-confidence model. The Y axis of the SHAP plot displays the top 30 most important features in descending order based on their feature importance in the model. The colour scale shows the values of the features (high or low, as indicated by the colour bar) and the X axis shows their corresponding impact on the model’s prediction (a positive value indicating the variant is more likely to be predicted as regulatory). (C) The comparison of the AUROC scores of human models trained on the entire set of general features, as indicated in Table 1, and human models trained on the common features available for both human and cattle, as highlighted in salmon colour in Table 1. The grey dashed line represents parity. The X and Y axis are limited from 0.70 to 1.0 for better clarity of the scatter points.

To understand which features impacted prediction outcomes, and therefore may be most relevant to training cattle models, a SHAP summary plot was generated for model interpretability (S. M. Lundberg & Lee, 2017). A SHAP summary plot visualizes the impact of different features on a machine learning model’s prediction, offering insights into feature importance and the direction of their influence (S. M. Lundberg & Lee, 2017). The SHAP values in the summary plot quantify each feature’s contribution to the model’s predictions, indicating both the magnitude and direction of their impact on individual data points. Figure 3B illustrates the top 30 most important features for the best-performing cross-tissue model, along with their corresponding impacts on the model’s predictions. Note the annotation categories described in Table 1 can be split across multiple features after feature encoding, such as the VEP variant effect predictions. The most influential feature is the distance to the promoter. As shown in the plot, a moderate distance-to-promoter value (around 0, as distance includes both positive and negative values), indicating proximity to the promoter region, is linked to a higher probability of a variant being predicted as a regulatory variant. Other features that prominently contribute to the model outcomes include gene density, variant consequence, distance to TSS, and conservation scores. In the tissue-specific models, in addition to features that exhibit importance across different models, such as gene density and distance to TSS, certain chromatin features emerged solely among the top features in specific tissue models. For example, the distance from the variant to open chromatin regions marked by H3K4me3 in the psoas muscle featured among the top 10 most important attributes in the muscle-specific model (Supplementary Fig. 6A). Similarly, the distance from the variant to open chromatin regions marked by H3K4me3 in the neutrophil cell line ranked among the top 5 most important features exclusively in the blood-specific model (Supplementary Fig. 6B). This suggests that tissue-specific chromatin data can, to some extent, improve tissue-specific model predictions.

### Comparison of human and cattle baseline models

There is a comparative lack of chromatin and other omics datasets for most livestock species compared to humans, but notably, many of the most important features in Figure 3B are readily available for non-human species. To examine this further we trained new human models, using the same set of human regulatory variants but excluding the human-specific omics datasets as features. This led to only modest drops in model performances (Figure 3C). While the human-specific features like distances to chromatin data and regulatory elements offer a slight boost to tissue-specific models (AUROC score increase of ∼0.03), their impact on cross-tissues models is minimal (only ∼0.007 improvement). Therefore, to directly compare the ability of predicting regulatory variants in different species, we trained cattle models on the same set of common features as highlighted by the salmon colour in Table 1. The cattle models were also evaluated using a chromosome-level holdout strategy (See Methods). Here, we present the model performance tested on chromosome 1 and 28 (chr1 and chr28). The key feature distributions of foreground and background variants in the training and test sets are shown in Supplementary Fig. 5. Results tested on other chromosomes can be found in Supplementary Fig. 7. The cross-tissue cattle model achieved a test AUROC score of 0.73 (Figure 4A), and a mean cross-validation AUROC score of 0.71 (Supplementary Fig. 7A). We repeated training and testing 10 times using different randomly selected matching background sets to assess the robustness of cattle cross-tissues model. The models yielded test AUROCs ranging from 0.71 to 0.74, with a mean test AUROC of 0.73 and a standard deviation of 0.01. Comparison of the feature importances for these human and cattle cross-tissue models trained on the common sets of over 470 features showed that the most important features determining the prediction results were largely consistent across the two models, with the top seven features being the same in both species (Figure 4B).

**Figure 4.**
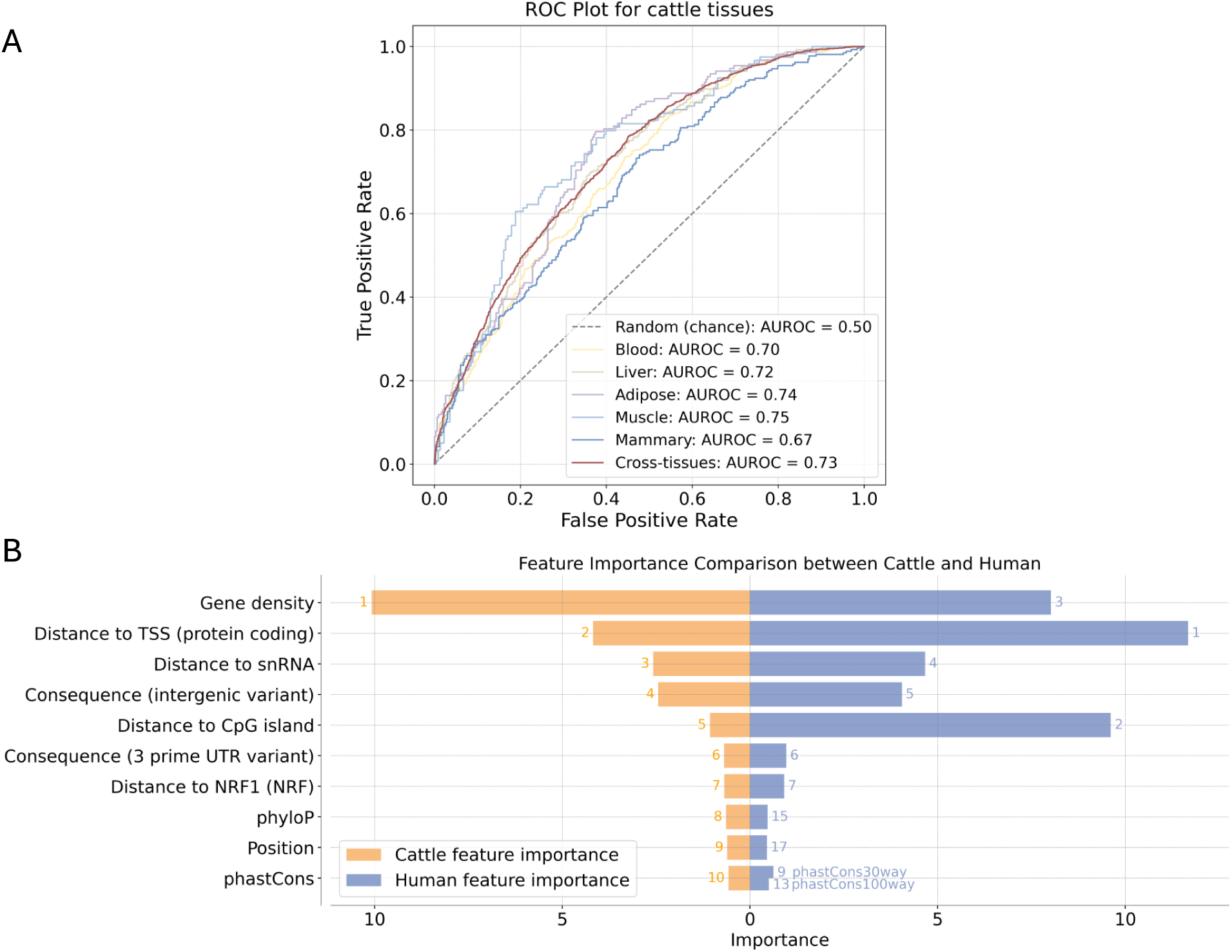
The comparison of cattle and human models trained on the common set of features. (A) ROC curves for cattle tissue-specific and cross-tissues models. The numbers in the plot legend represent the AUROC scores of different models. (B) Bar plot showing the top features in the cattle (orange) and human (blue) cross-tissues models trained on the common features. The X axis represents the relative feature importance in cattle and human models, respectively. The Y axis shows the top 10 features in the cattle model. The numbers next to the bars represent the rank of each feature. In humans, two phastCons conservation scores were available (phastCons30way and phastCons100way), ranking 9^th^ and 13^th^.

Tissue-specific regulatory variants for adipose, liver, blood, mammary, and muscle were also extracted for cattle. These sets comprised 6,308, 13,034, 16,328, 10,376, and 5,378 variants, respectively (each set includes both regulatory and background variants). Among cattle tissue-specific models, the muscle model achieved the highest test AUROC score of 0.75 (Figure 4A) (mean cross-validation AUROC score of 0.76). In general, cattle models achieved poorer performance compared to human models when trained on the equivalent set of features.

To explore whether incorporating cattle chromatin data could improve model performance, we included 95 cattle chromatin datasets. The features were used both in a cross-tissue manner and in a tissue-specific manner for available tissues (liver, adipose, and muscle). However, the model’s performance remained the same (Supplementary Fig. 8), suggesting the addition of the cattle chromatin data had little improvement on the accuracy of predicting regulatory variants.

Given the difference in the number of human and cattle variants available for training, we down-sampled the human variant sets to match the number of variants in cattle models for a fair comparison. Despite this adjustment, the human models still performed better (Supplementary Fig. 9). These observations suggest that the performance gap between species is potentially largely driven by differences in the quality of the regulatory variant sets used, rather than limitations in the available feature sets.

### Human omics data does not improve cattle models

Previous studies have shown deep conservation of some regulatory elements in animals (Wong et al., 2020), potentially enabling annotation transfer between related species like humans and cattle. To explore this, we characterized the ability of using human annotations to improve cattle regulatory variant prediction. Three approaches were tested: applying pre-trained human models to cattle variants, applying incremental learning based on the human model for cattle prediction, and lifting cattle variants to the human genome to train the model using human annotations. Note that in the latter only about 52% of the cattle GTEx variants could be lifted over to the human genome. These approaches achieved test AUROC scores of 0.67, 0.68, and 0.68, respectively, which are lower than those of the cattle models trained on all cattle annotations except for cattle chromatin data (AUROC = 0.73). Therefore, this outcome highlights that species-specific models outperform, even when fewer annotations are available. Human-based models, while capturing some general principles, may miss species-specific intricacies crucial for accurate predictions.

### Enformer provides little benefit in predicting regulatory variants in either species

Enformer is a sequence-based model that can predict a range of regulatory features from sequence alone, and the authors illustrated its potential at predicting regulatory variants by comparing regulatory feature predictions when variant alleles are changed in the input sequence (Avsec et al., 2021). Such models therefore provide a potential avenue to enhance feature sets from just genome sequences, and if it could, at least in part, be applied across species, it would have the potential to fill in gaps in omics data for less well characterised mammals. We consequently investigated the usefulness of features derived from the sequence-based Enformer model to inform both human and cattle predictions. The human high-confidence cross-tissues dataset was annotated with 5,313 Enformer features, i.e. the difference in predicted omics data between genomic sequences with reference and alternative alleles. We then trained CatBoost models using three feature sets: Enformer features only, general features only, and a combination of both. As shown in Figure 5A, the model trained with general features performed slightly better than the one using only Enformer features (AUROC score increase of 0.02). The combined model achieved a slight improvement in performance, with an AUROC score increase of 0.01 compared to the general features model. This suggests that while incorporating Enformer features may provide some limited additional regulatory information, its overall contribution to predictive power remains modest. We further applied the Enformer sub-workflow to annotate cattle tissue-specific variants. This aimed to explore whether these inferred chromatin features could improve cattle model predictions, given the comparative lack of omics data for this species. We initially applied the full set of Enformer features but the performance remained unchanged compared to not using Enformer features (Supplementary Fig. 10). Subsequently we incorporated tissue-relevant Enformer features into the cattle tissue-specific models, which resulted in a modest improvement in performance, with a maximum AUROC score increase of 0.005 (Figure 5B). Our findings suggest that Enformer features offer limited improvement to regulatory variant prediction model performance in either species. These results are broadly consistent with recent work by Huang et al. highlighting limitations in some deep learning models’ ability to capture the intricacies of individual genetic changes on gene expression regulation, especially those at distal sites (Huang et al., 2023).

**Figure 5.**
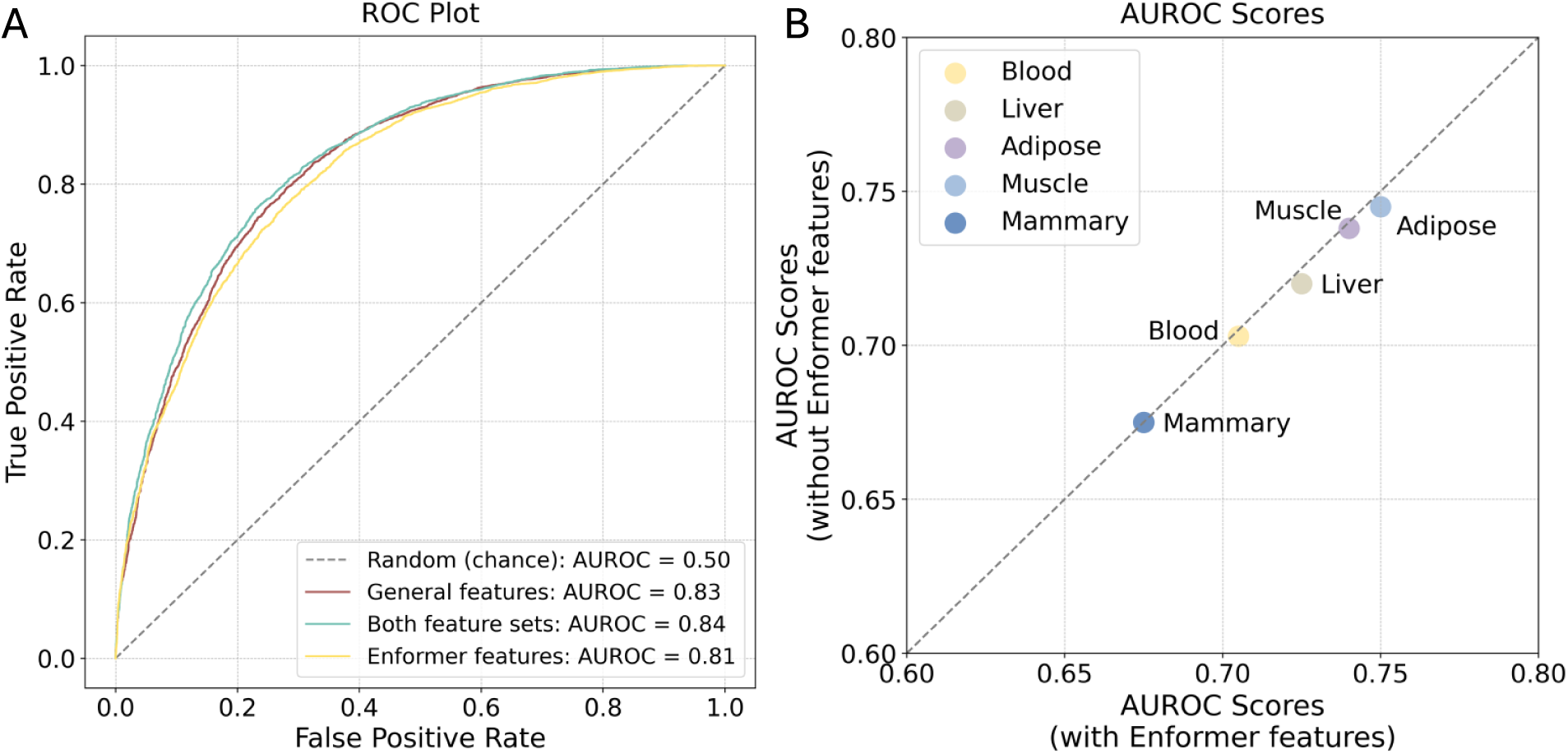
The comparison of human and cattle models trained with or without Enformer inferred features. (A) ROC curves for human high-confidence cross-tissues model trained using Enformer features, general features, and the combination of two feature sets. The numbers in the plot represent the AUROC scores of different models. (B) The comparison of the AUROC scores of cattle tissue-specific models trained with or without tissue-specific Enformer features. The grey dashed line represents parity. The X and Y axis are limited from 0.60 to 0.80 for better clarity of the scatter plot.

### Prediction performance at distal regulatory elements

To evaluate the effectiveness of our models at predicting regulatory variants across diverse genomic regions, we compared their performance on variants stratified into five distance ranges relative to TSSs in different tissues. For each distance bin, the major class (i.e., the class with more variants) was down-sampled to match the number of the minor class, creating a balanced set. Bins were defined to ensure comparable variant counts after down-sampling within each tissue. In cattle, tissues with fewer than 10,000 variants were excluded. As illustrated in Figure 6, model performance displayed a slight downward trend with increasing distance from the TSS. Despite this, the overall stability of performance, even for distal variants, suggests that our model generalizes effectively across the genome, including regions far from canonical regulatory hubs.

**Figure 6.**
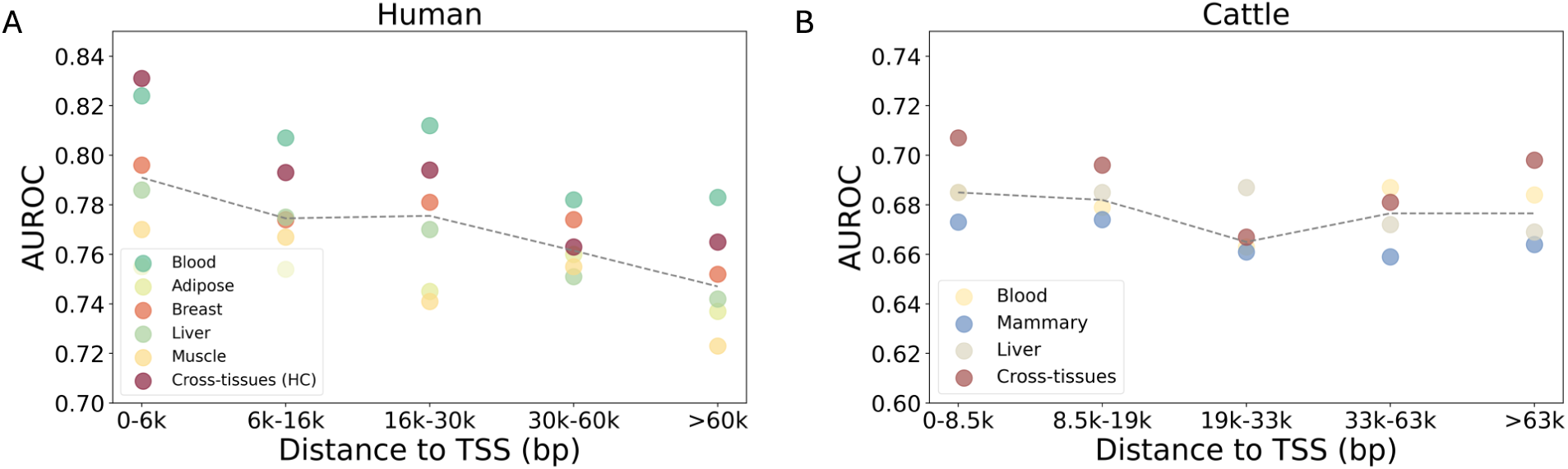
Human and cattle model performance using variants within different distance ranges to the TSS. (A) Human tissue-specific and cross-tissues high-confidence (HC) models performance based on variants in different TSS bins. The grey dashed line is plotted based on the median test AUROC score among the six datasets in each range. (B) Cattle tissue-specific and cross-tissues models performance based on variants in different TSS bins. The adipose and muscle datasets were excluded due to their relatively low variant counts. The grey dashed line is plotted based on the median test AUROC score among the four datasets in each range.

### Model performance in predicting human and cattle MPRA regulatory variants

eQTLs have several limitations when used to train machine learning models. Most notably, only a subset of variants, such as those found above a certain allele frequency and within a certain distance of genes (i.e. cis-eQTLs), are tested when characterising eQTLs. Previous work has shown that the distribution of eQTLs in the genome is biased towards genes under less constraint and fewer functional annotations compared to variants linked to diseases and traits (Mostafavi et al., 2023). Precisely mapping the functional variant at a locus is also difficult, even when using statistical fine-mapping approaches, suggesting that eQTLs do not fully capture the spectrum of regulatory variants in the genome.

Massively parallel reporter assays (MPRA) are an alternative approach for mapping regulatory variation in the genome that overcome some of these limitations. For example, approaches such as SuRE can test largely every heterozygote variant in the genome irrespective of its population-wise allele frequency or distance to known genes (van Arensbergen et al., 2019). Likewise, for most variants the confounder of LD is removed, as each variant is tested independently, allowing for the effect of individual variants on transcription to be characterised. MPRA approaches do though have their own limitations, most notably the fact that the variants are tested outside of their native sequence and chromatin environment. Likewise, they can only be carried out in suitably transfectable cell lines. These methods are therefore complementary approaches for defining regulatory variation.

A further advantage of examining SuRE data is that the data is potentially more comparable between species, having been generated using the exact same approach (SURE), though in different cell types. This allowed us to further explore whether differences in model performances between species was due to differences in the training sets of regulatory variants. Consequently, in addition to using regulatory variants from the GTEx project, we also built the models using regulatory variants (raQTLs) based on the SuRE technique. For our human SuRE models, we included regulatory variants identified in two cell lines: K562 (a myelogenous leukemia cell line) and HepG2 (a liver hepatocellular carcinoma cell line) (van Arensbergen et al., 2019). We also trained a model on the combined dataset of these two lines. These three human SuRE models achieved test AUROC scores of 0.82 (HepG2), 0.84 (K562), and 0.81 (combined dataset) on their respective test sets. Their ROC curves and cross-validation results are shown in Supplementary Fig. 11. Additionally, we identified the top 30 most important features in raQTL models (Supplementary Fig. 12). Among all the features, distance to motifs, distance to cell type-specific chromatin data, and conservation score were consistently ranked among the top features in both cell lines. While features such as distance to TSS and gene density, which are among the top features in the GTEx models, did not appear in the top-ranked features of raQTL models. This difference likely reflects the distinct regulatory landscapes captured by these models. SuRE assays measure the intrinsic regulatory potential of DNA sequences in a standardized context and regulatory environment independent manner. As a result, features related to intrinsic sequence properties, such as distance to motifs and conservation scores, rank higher in raQTL models.

For cattle modelling, we used a set of 66,501 SuRE raQTLs recently derived from cattle aortic endothelial cells (unpublished). To explore the potential optimal p-value threshold for defining raQTLs in cattle, we applied different FDR adjusted p-value thresholds. Equivalent numbers of background variants were randomly down-sampled from the tested variants with adjusted p-values greater than 0.05. As shown in Figure 7, as the p-value threshold decreases, the AUROC score plateaus at around 0.81, broadly comparable to that observed for the human models. However, noise becomes apparent when the p-value threshold is smaller than 1 × 10^−10^, mainly due to the fewer variants included in the training and testing sets when below this cutoff. We compared the top 30 most important features in the cattle raQTL model with the human raQTL models and found that distance to motifs and conservation scores ranked among the top features in both species, as shown in Supplementary Fig 12. Consequently, the difference between the human and cattle models is not apparent here when trained on more comparable datasets of regulatory variants.

**Figure 7.**
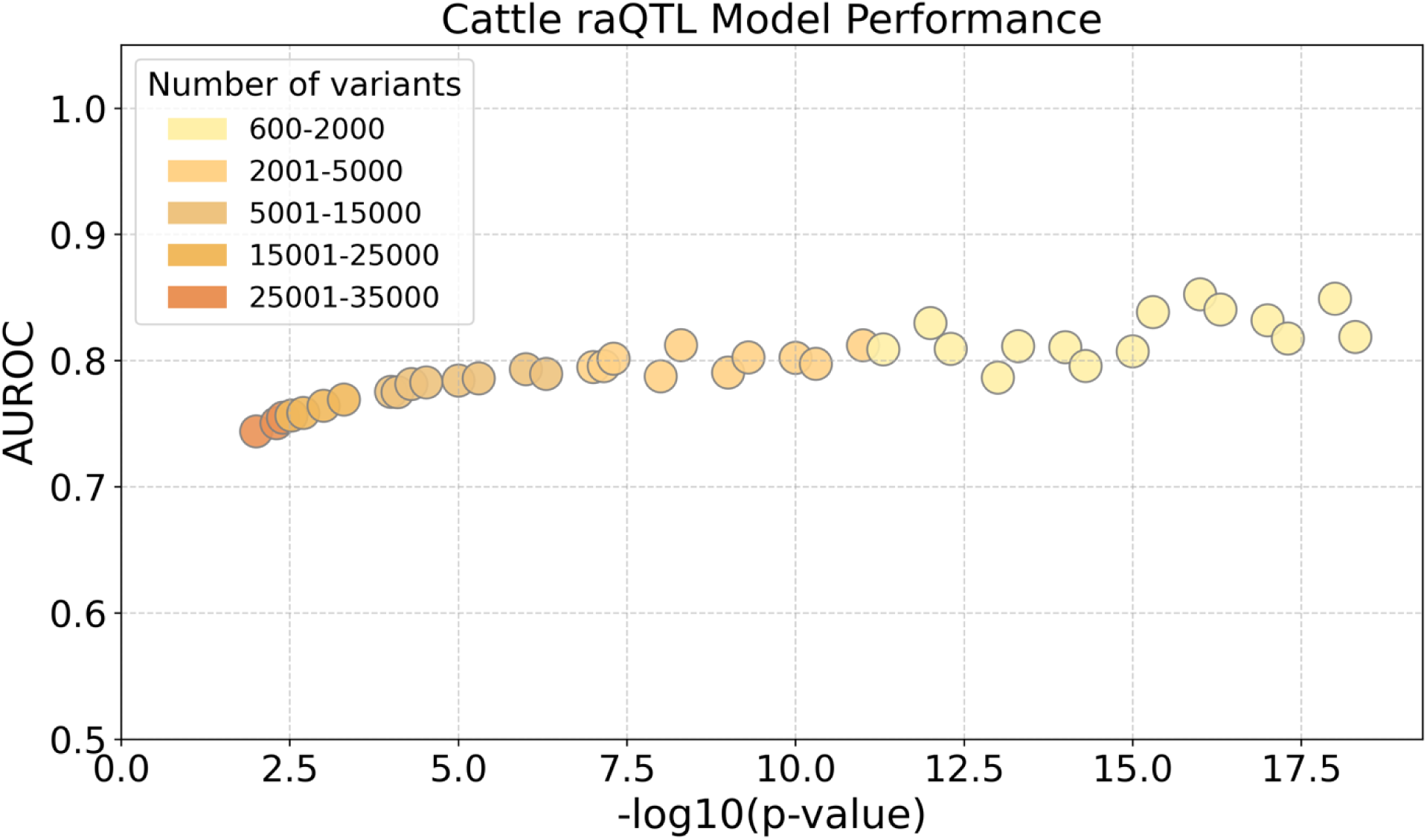
Cattle raQTL model performance when training and testing on regulatory variants selected based on different FDR adjusted p-value thresholds. For better clarity, the X axis is transformed to 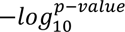. The colour of each point represents the number of raQTL variants above the given threshold.

### Our models substantially enrich for trait-associated variants in both species

A key motivation for training models that can predict regulatory variants is to identify which variants at loci linked to diseases and traits may be the functional drivers of the phenotypes. Likewise simply being able to enrich for functional variants among the set of genetic changes used to calculate genomic estimated breeding values is expected to improve their accuracy (Xiang et al., 2021). We therefore investigated the generalizability of our human and cattle regulatory models to predict functional GWAS variants. Of 28,918 human GWAS lead variants obtained from the credible sets analysis of the Open Target Genetics project (see Method), approximately 75% were identified as regulatory with a prediction probability exceeding 0.5 using the cross-tissues high-confidence human GTEx model. This is an enrichment ratio of 2.3 compared to a background dataset of the same size, with randomly selected variants (repeated 10 times), where an average of 32% were predicted as regulatory at the same threshold (standard error of the mean: 0.62%).

The marked shift in prediction probability distributions between putative functional and background variants is shown in Figure 8A. To explore the impact of the prediction probability threshold for distinguishing regulatory variants in the model, different thresholds were utilised, and the corresponding ratio of the proportion of lead to background variants predicted as being regulatory was recalculated. As shown in Figure 8B, it is possible to observe over 10-fold enrichments for functional variants at higher thresholds, suggesting these models can substantially enrich for variants driving downstream phenotypes.

**Figure 8.**
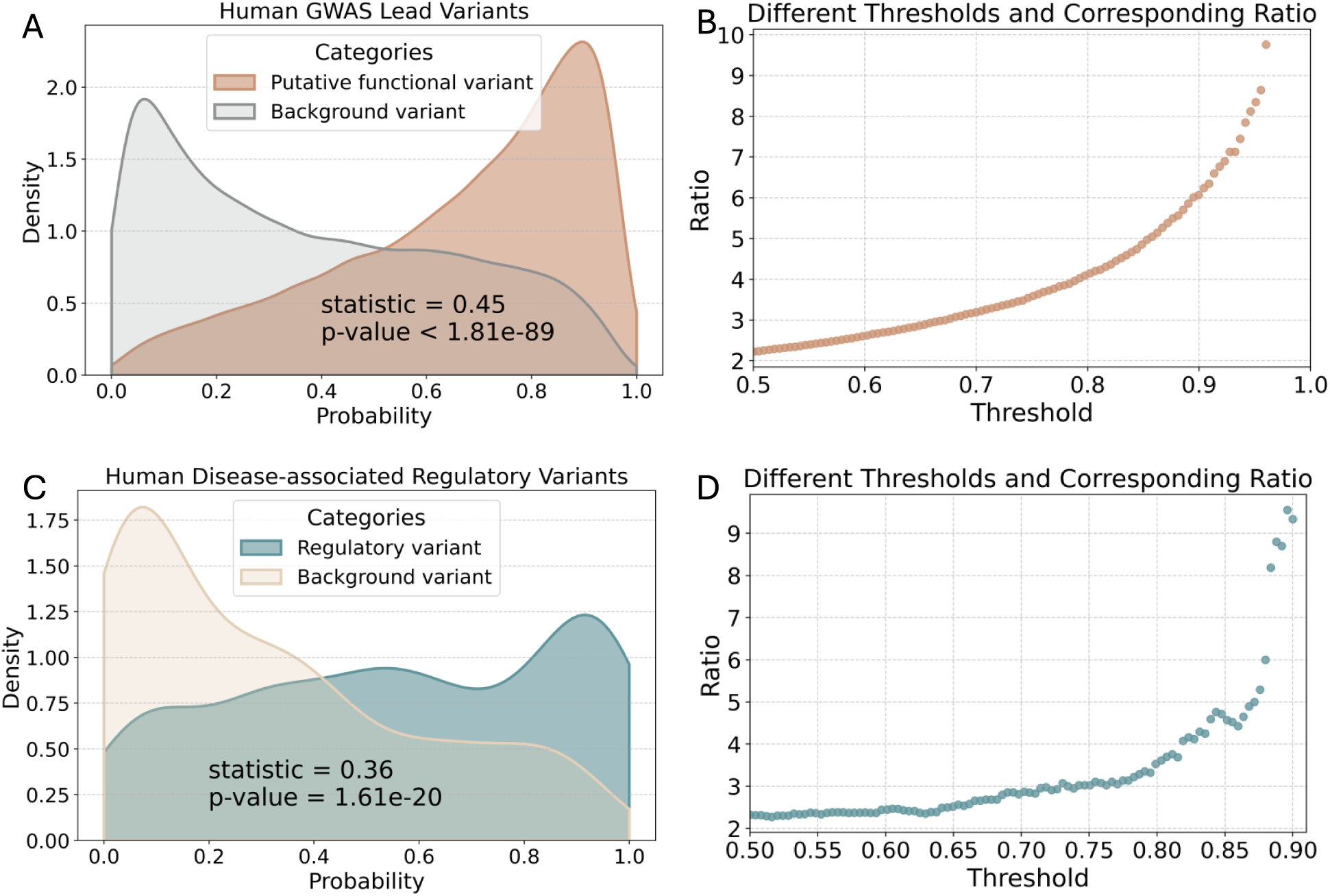
Prioritising human functional variants. (A) Distributions of model derived prediction probabilities of putative human GWAS functional variants and background variants. The Kolmogorov-Smirnov statistic and p-value are shown in the plot. (B) Different prediction probability thresholds and their corresponding ratios of the proportion of GWAS lead variants predicted as regulatory to the proportion of background variants predicted as regulatory. (C) Distributions of model derived prediction probabilities of human disease-associated regulatory variants and background variants. The Kolmogorov-Smirnov statistic and p-value are shown in the plot. (D) Different prediction probability thresholds and their corresponding ratios of the proportion of human disease-associated regulatory variants predicted as regulatory to the proportion of background variants predicted as regulatory.

To further assess the high-confidence human model’s ability at predicting the regulatory effect of variants, we obtained 406 non-coding Mendelian disease-associated regulatory mutations (Smedley et al., 2016). After lifting these variants from GRCh37 to GRCh38, 345 single nucleotide variants on autosomes were retained for analysis. Notably, at a prediction probability cutoff of 0.5, 58.0% of these validated regulatory variants were predicted as regulatory, compared to an average of 22.9% of matching background variants randomly selected from the 1000 Genomes cohorts (repeated 10 times, standard error of the mean: 0.71%). Figure 8C illustrates the difference in the prediction probability distributions between these groups and Figure 8D explores how different prediction probability threshold impacts the ratios of regulatory variants predicted as regulatory to the background proportion, with ratios of up to 9-fold observed. The high proportion of both GWAS and disease-associated variants predicted as regulatory by the model demonstrates the generalizability and effectiveness of our approach in prioritizing regulatory variants linked to downstream phenotypes.

To explore the utility of our cattle models at prioritizing variants linked to important cattle phenotypes we obtained GWAS data for four cattle traits: body weight, stature, body condition score (Reynolds et al., 2021), and milk yield (Jiang et al., 2019). Using GCTA-COJO, we selected independently associated SNPs for body weight, stature, and body condition score based on the corresponding GWAS summary statistics data. Specifically, 344, 241, and 96 fine-mapped SNPs were selected from the conditional analyses respectively. Additionally, we obtained 210 fine-mapped variants for milk yield from Jiang et al.. Table 2 summarizes the proportion of these variants predicted as regulatory using our different cattle models. All models predicted a higher proportion of fine-mapped variants as regulatory compared to the equivalent number of matching background variants. At the default prediction probability threshold of 0.5, the ratios of the proportion of fine-mapped GWAS variants predicted as regulatory to the proportion of background variants predicted as regulatory ranged from 1.5 to 3.0. Figure 9 A,B,C illustrate the prediction probability distribution differences between fine-mapped and background data in the three traits, body weight, body condition score, and milk yield. The prediction probability distribution plots for stature can be found in Supplementary Fig. 13. We further explored the impact of different prediction probability thresholds in the cross-species model on the ratios of the proportion of fine-mapped variants predicted as regulatory to background proportion. As shown in Figure 9D and E, 10-fold and 18-fold enrichment ratios are observed at higher thresholds for body condition and milk yield, respectively. This suggests that a higher prediction probability threshold could be useful for identifying high-confidence functional variants in GWAS datasets (at the cost of a higher false negative rate).

**Table 2.**
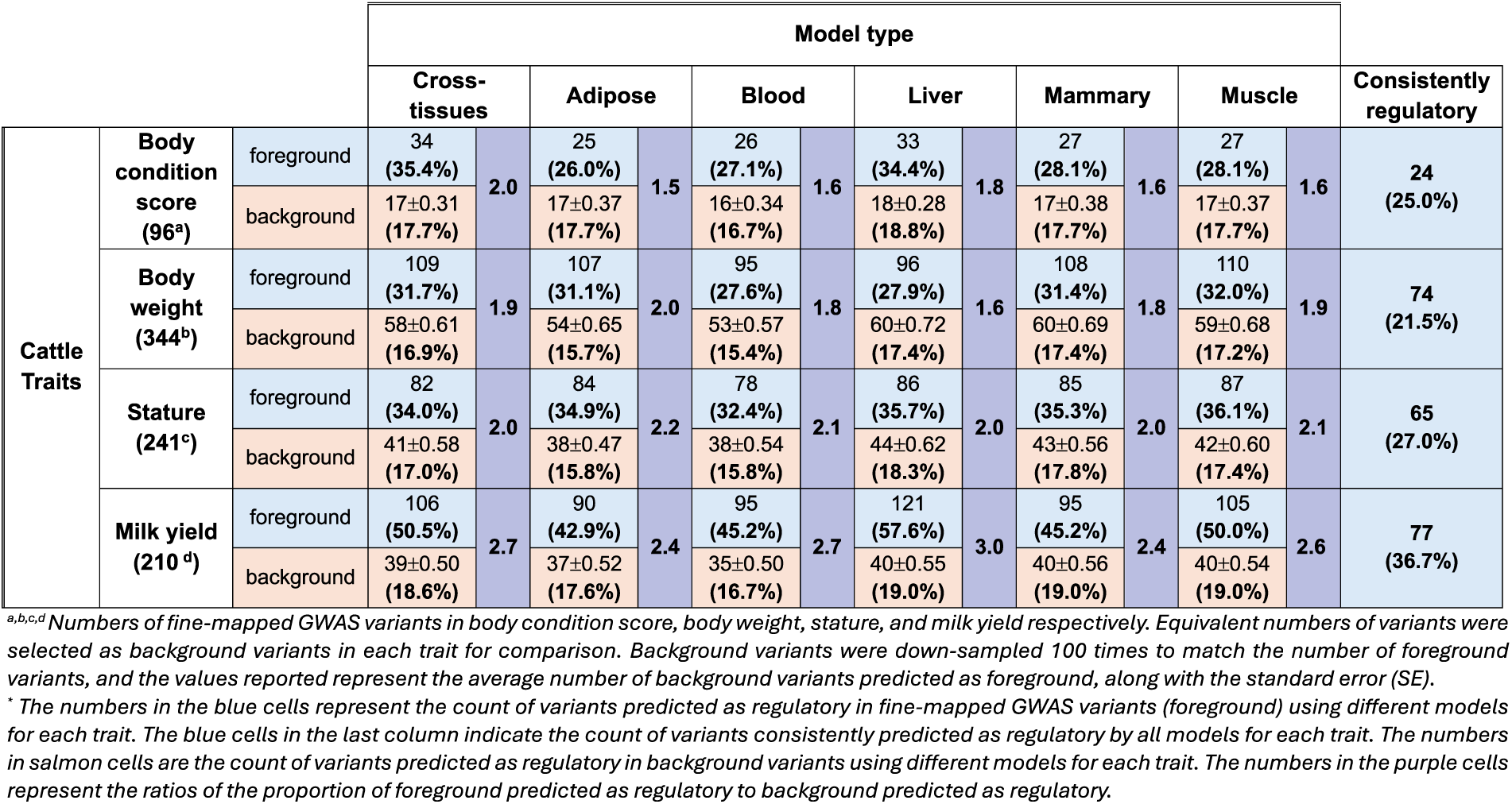
The proportion of variants predicted as regulatory in four cattle traits using different models.

**Figure 9.**
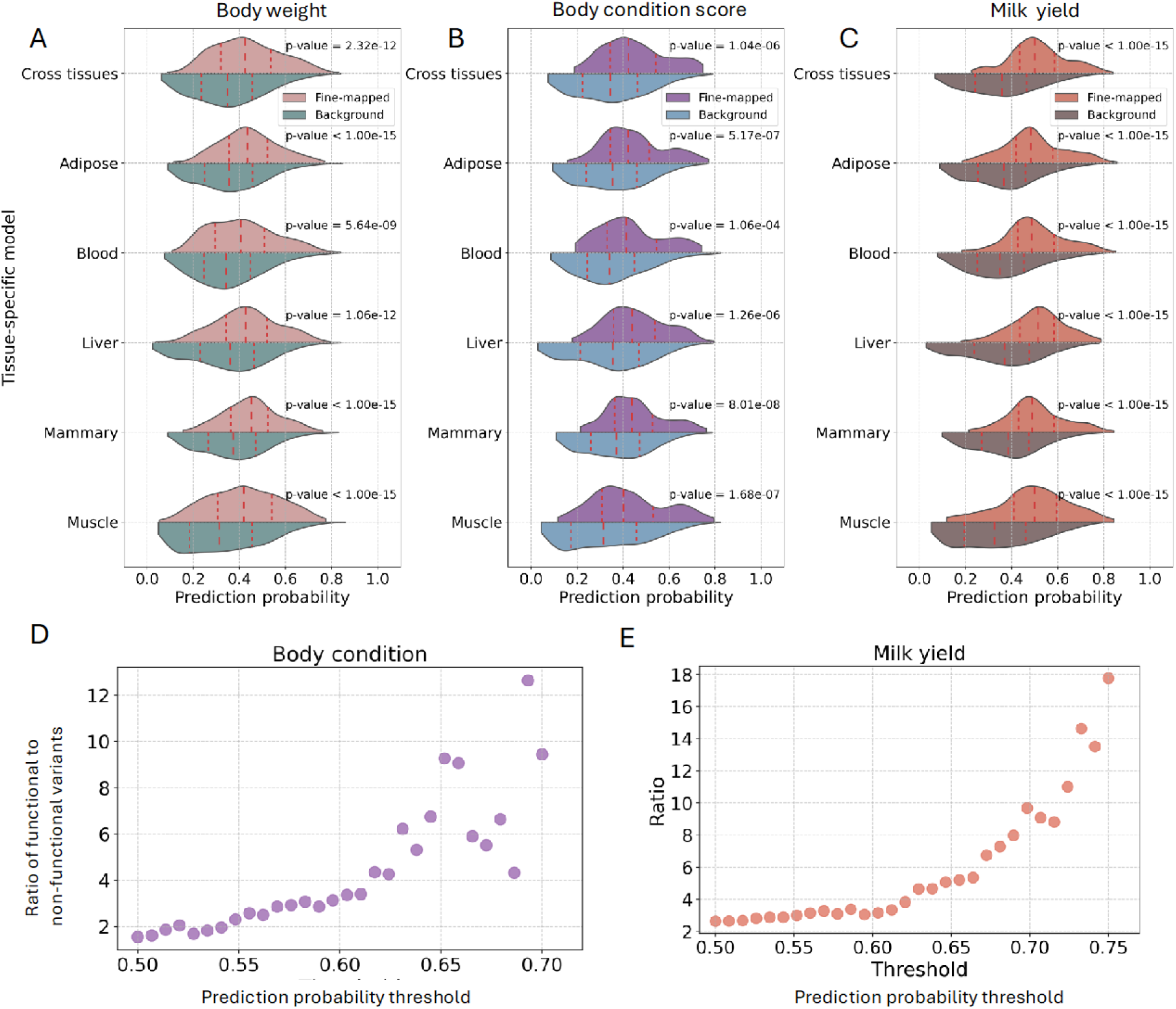
Prioritising cattle functional variants. Distributions of model derived prediction probabilities of cattle GWAS fine-mapped and background variants for (A) body weight, (B) body condition, and (C) milk yield. The red dashed lines in the violin plots represent the median and quartile values. All prediction probabilities are significantly different between fine-mapped and background groups in each experiment. Their corresponding p-values are displayed in the figure (Two-sample Kolmogorov-Smirnov test). (D) Different prediction probability thresholds and their corresponding ratios of the proportion of cattle GWAS fine-mapped variants associated with body condition score predicted as regulatory to the proportion of background variants predicted as regulatory using the cross-tissues model. (E) Different prediction probability thresholds and their corresponding ratios of the proportion of cattle GWAS lead variants associated with milk yield predicted as regulatory to the proportion of background variants predicted as regulatory using the cross-tissues model.

We identified variants that were consistently predicted as regulatory across different cattle models. Among the 96 body condition score variants, 24 of them were predicted as regulatory by all six models with a prediction probability exceeding 0.5. Similarly, 74 out of 344 body weight-associated SNPs, 65 out of 241 stature-associated SNPs, and 77 out of 210 milk yield-associated SNPs were consistently predicted as regulatory by all six models with a prediction probability greater than 0.5 (Table 2). Supplementary Fig. 14 illustrates three examples. The chr14:23070083_C/T SNP, located near the TSS region of the *TGS1* gene, was predicted as a regulatory variant associated with stature (Supplementary Fig. 14A). This gene has been found to be associated with stature in both human and cattle populations (Zhao et al., 2015). The body weight-associated variant chr5:103942835_G/A is found in the intergenic region upstream of the *CD27* gene (Supplementary Fig. 14B), which has previously been associated with weight gain in mice (Englebert et al., 2024). Another predicted body condition score associated variant, chr2:125968341_G/A, was found near the *WDTC1* gene (Supplementary Fig. 14C). This gene has been associated with fat accumulation and obesity in humans (Ducos et al., 2017).

### Predicting regulatory elements linked to disease

These models effectively predict at which genomic locations a DNA change is expected to have a regulatory effect. Consequently, their utility may be broader than just predicting regulatory variants but also the location of key regulatory elements in the genome. One of the few examples in cattle where a distal regulatory element has been directly linked to a downstream phenotype, is an enhancer element downstream of the GC gene. In a previous study (Lee et al., 2021), disruption of this element by a copy number variant was linked to multiple traits including mastitis resistance, body condition score and milk production traits. Figure 10 A-C shows this locus and the blood model prediction probabilities for variants in this region when trained just on the baseline features marked in the salmon colour in Table 1. The variant with the highest prediction probability was chr6:86954490_C/T, that was not only the lead variant in the body condition score GWAS but also located at the enhancer region of the GC gene linked to the downstream phenotypes. So, despite the model not being trained using chromatin or GWAS data, it successfully identified the lead variant at this locus at the enhancer regulatory region. Together these results indicate the cattle models may have possible potential for prioritising regulatory variants and elements linked to important cattle traits that could be explored in further work.

**Figure 10.**
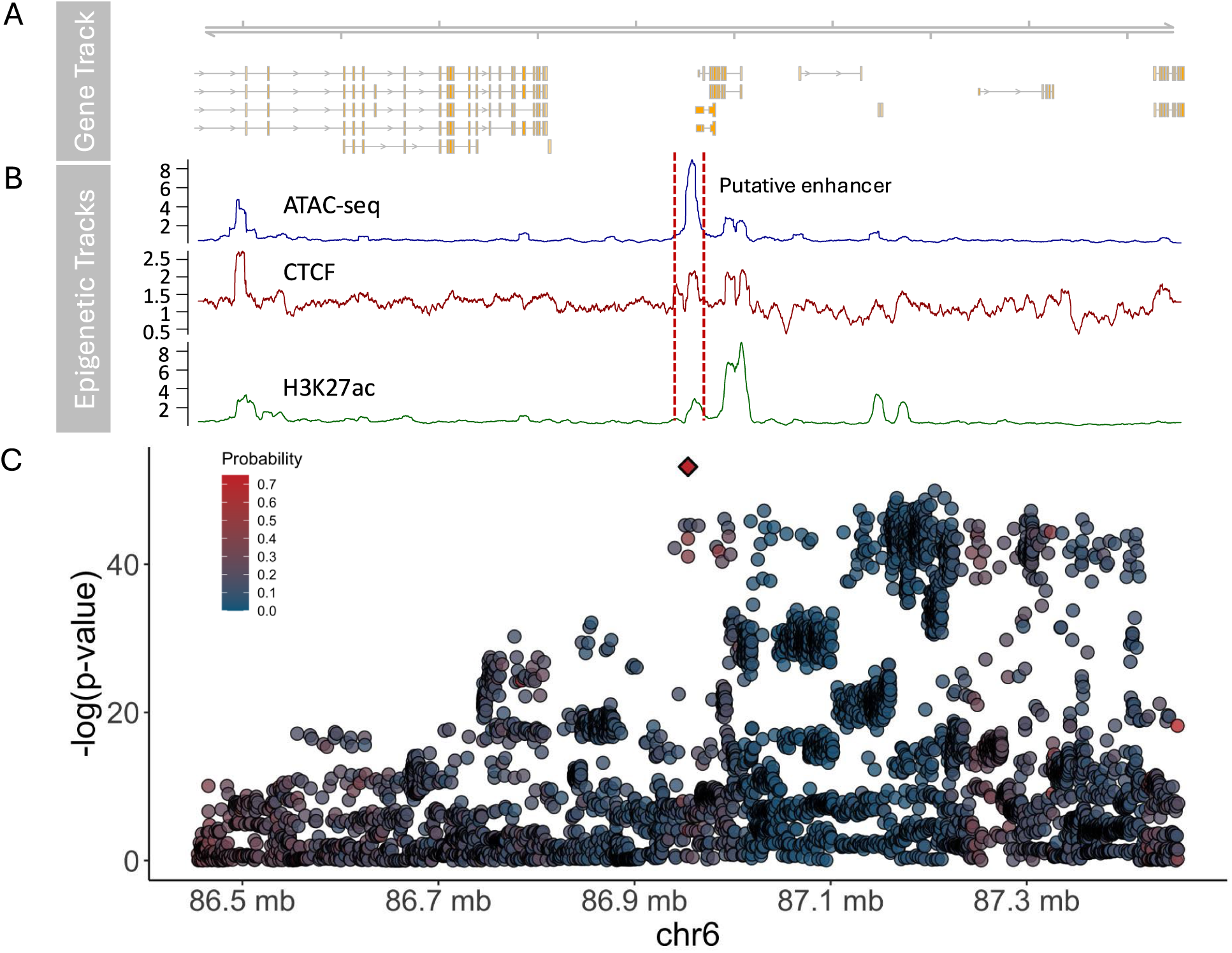
Regulatory predictions at the GC locus linked to traits including mastitis resistance and body condition score. (A) The genes and different isoforms at this locus. (B) ATAC-seq (Lee et al., 2021), CTCF and H3K27ac (Kern et al., 2021) tracks in this region, with the location of the putative trait-associated enhancer identified by Lee et al. indicated by the dashed lines. (C) Strength of association of cattle variants in the region with body condition score (Y axis), against their position (X axis). Colour indicates the prediction probabilities of each variant from the cattle blood model. Variant chr6:86954490_C/T is represented by a diamond.

## Discussion

Identifying functional regulatory variants remains a considerable challenge, in large part because they typically occur in less well-annotated non-coding regions and are affected by linkage disequilibrium (LD) among neighbouring variants. These problems are even more pronounced in livestock species, where high-quality functional genomic data is relatively scarce and LD tends to be extensive. To address this challenge, we developed a species-agnostic variant annotation pipeline that integrates a range of features, from sequence conservation scores to positional properties, and applied it to build human and cattle machine learning models for predicting variants with regulatory effects.

Our models, trained in both humans and cattle, effectively distinguished regulatory variants from non-regulatory variants, and similar predictive features emerged as important in both species. Earlier work on prioritising cattle functional variants had highlighted the particular value of conservation metrics (Xiang et al., 2019). However, that study had relied on lifting over human conservation metrics to the cattle genome which is problematic because only about half of human bases can be directly mapped to a corresponding cattle base. In the present study, we generated PhastCons and PhyloP conservation metrics specifically for cattle for the first time, confirming their importance for predicting regulatory variants in line with previous findings. Although the multiple alignment underlying these scores was based on the Btau_5.0.1 reference genome, 98.7% of the metric sites successfully lifted over to BosTau9, illustrating that the new cattle-specific conservation metrics can be effectively utilised with current genome builds.

The performance of both the human and cattle models exhibited a slight downward trend in performance with increasing distance from TSSs. This pattern is perhaps expected given the more defined regulatory code near promoters and other well-characterized transcriptional control regions. However, despite this modest decline, the overall stability of performance across all distance bins suggests that our models have the ability to generalize across the genome, including regions far from canonical regulatory hubs. Interestingly the NRF1 motif was found to be the most informative motif for predicting regulatory variants and the seventh most important feature overall in both species when training was restricted to the set of common features. This may result from NRF1 being a ubiquitously expressed canonical activator with a relatively well defined motif (Duttke et al., 2024).

Applying our GTEx-based models to human and cattle datasets demonstrates their effectiveness in prioritizing functional variants associated with important traits. While the precise circumstances and full extent of improvements remain a topic of ongoing research, previous studies suggest that incorporating functional variant-based SNP panels may enhance trait prediction accuracy (Santana et al., 2023; Xiang et al., 2021). Consequently the enrichments of up to 18-fold for trait-associated variants holds potential for the prioritization of variants for calculating genomic estimated breeding values (gEBVs) in livestock. Unlike the requirement for the direct identification of functional variants to improve livestock using approaches such as genome editing, the identification of causal variants is not always required when prioritizing SNPs for gEBV calculation, provided that the variants in LD with the true causal alleles are enriched for (Xiang et al., 2021). Consequently, perfect classification accuracy may not be strictly necessary to achieve practical gains. Simply enriching the SNP panel used for gEBV calculations with functional variants, or tagging variants in high LD, has the possibility to improve their accuracies, and potentially lead to substantial cumulative benefits over time.

The analysis of the GC locus also highlights the possible potential of these models for fine-mapping functional regulatory elements at GWAS loci. Despite not being trained on the GWAS or chromatin data, the model identified the lead GWAS variant at the previously identified enhancer element as the variant with the highest regulatory potential. So, although the functional variant at this locus is thought to be a structural variant, this result highlights how relevant regulatory elements may potentially be prioritised using these models for further follow up study.

To date, deep learning-based models, especially those popular models in natural language processing (NLP), have been explored for understanding DNA sequences. However, the prediction results in our study illustrate that Enformer features did not provide additional substantial discriminative information beyond our general features for both human and cattle models. This aligns with recent studies evaluating popular deep learning architectures, including Enformer, for predicting gene expression variation (Huang et al., 2023), as well as *in silico* investigations focusing on Enformer’s ability to capture expression in promoters and enhancers (Karollus et al., 2023). These observations reported limitations in these models’ performance. Specifically, they observed limited effectiveness in elucidating the impact of variants on gene expression and difficulties in capturing the causal influence of distal enhancers on gene expression. Although these previous studies primarily focused on Enformer’s ability to predict gene expression levels, rather than simply define the location of regulatory variants, these observations regarding Enformer’s limitations align with the results of our study, highlighting the challenges deep learning models face in capturing the complex regulatory effects of individual base changes.

The difference in performance between human and cattle models, even when using common features, suggests limitations in the cattle variant data. This is further supported by the minimal improvement (∼0.03 AUROC score increase) observed in human models trained on all general features compared to models using only common human-cattle features, and by the fact that adding 95 cattle chromatin features did not enhance the performance of the cattle model. The comparatively small gap in performance of the cattle and human models trained on the more comparable MPRA data also supports the idea that the difference in performance of the cattle and human eQTL models is because the foreground sets are less well matched, i.e. the core issue lies in the lack of a gold-standard regulatory variant set for training cattle models, which is largely responsible for their relatively low performance. This limitation is likely exacerbated by strong linkage disequilibrium (LD) in cattle populations, which complicates fine-mapping efforts from eQTL data and may introduce uncertainty in regulatory variant identification. While the cattle GTEx project serves as one of the most extensive reference resources for the cattle transcriptome available to date, it does have challenges. This includes a comparatively limited number of individuals and breeds for certain cell types, the dependence on calling variants from the RNA-seq data, and difficulty of accurate variant imputation in underrepresented breeds. Additionally, unlike human GTEx, cattle GTEx data comes from diverse studies, potentially reducing its reliability due to variable batch effects. Together these factors may be leading to the difference in model performance between human and cattle. Addressing these challenges requires larger sample sizes in consistent cohorts, matching whole-genome variant calls, cross-breed analyses to break down breed-specific LD patterns, and potentially improved functional annotation efforts using high-throughput cellular screens. A more comprehensive collection of high-quality cattle data, as well as data for other species, encompassing high quality functional data would be indispensable. With the increased availability of high-quality training data expected to further improve the performance of livestock models in prioritizing novel functional variants, and ultimately improving advanced breeding approaches.

## Methods

### Variant annotation pipeline

We obtained a range of annotations for variants from different sources, including sequence conservation, variant position properties, VEP annotations, sequence context, and predicted functional genomic data based on Enformer. Bigwig files for four different human conservation scores (phastCons100way, phastCons30way, phyloP100way, phyloP30way) were downloaded from UCSC (Nassar et al., 2023). The Python package pyBigWig (Ramírez et al., 2016) was then used to extract the corresponding score for each variant from the bigwig files. For cattle, phastCons and phyloP scores were not publicly available. Therefore, the 241-way mammalian alignment from the Zoonomia project was used to calculate the conservation scores for cattle (Armstrong et al., 2020). The hierarchical alignment (HAL) format 241-way multiple alignment was first converted to multiple alignment format (MAF) by chromosome using the hal2maf tool from HAL toolkit (Hickey et al., 2013), with the cattle genome serving as the reference. Meanwhile, the general feature format (GFF) file for the corresponding cattle genome (Btau_5.0.1) included in the Cactus alignment was downloaded from NCBI (Sayers et al., 2022), and the coding sequences (CDS) were extracted from the GFF files. Then the CDS in bed format together with the MAF files for each chromosome were used as inputs to the msa_view tool from the PHAST package (Hubisz et al., 2011) to generate four-fold degenerate sites (4d sites) in sufficient statistics (ss) format. The combined 4d sites file for the whole cattle genome was then randomly down-sampled to 40% of their original size to improve the computational efficiency in the downstream analysis. The down-sampled 4d sites and the phylogenetic tree of the HAL alignment were used as inputs to the phyloFit command (Hubisz et al., 2011) to estimate the neutral model. Subsequently, the conservation scores for cattle were computed using the phastCons and phyloP tools (Hubisz et al., 2011) with the neutral model serving as the phylogenetic model. Since the 241-way mammalian alignment employed the Btau_5.0.1 cattle genome assembly, we conducted a lift-over step to map the conservation scores to the updated BosTau9 assembly (Rosen et al., 2020). Due to the lack of lift-over chain file between these two assemblies, nf-LO (Talenti & Prendergast, 2021) was used to generate the chain file. Subsequently, the USCS liftOver tool was used to perform the mapping, successfully aligning 98.7% of variants to the BosTau9 cattle assembly.

For the variant position properties, distances from the variants to various genomic elements were calculated. The genomic locations of CpG islands for different species were downloaded from UCSC (Nassar et al., 2023). A total of 1,554 distinct types of human chromatin data were obtained from Ensembl (Martin et al., 2023). For cattle, 95 chromatin data features were obtained (Kern et al., 2021). Distances from the variants to the nearest CpG island and chromatin region were calculated using the bedtools closest function (Quinlan & Hall, 2010). The R library AnnotationHub was used to obtain transcription start sites (TSS) for different species, and only biotypes with greater than 1000 TSS were retained. The human regulatory features, including enhancers, transcription factors (TF), and CTCF binding sites, were obtained from Ensembl using the R library biomaRt (Smedley et al., 2009). Distances to the nearest transcription start site (TSS) and regulatory features (human only) were calculated using the ChIPpeakAnno R library (Zhu et al., 2010). Gene density per megabase pair (1Mb) for different species was calculated using the AnnotationHub R library (Gentleman et al., 2004) and GenomicRanges library (Lawrence et al., 2013). The VEP command line tool (McLaren et al., 2016) was used to obtain variant consequences. To capture the sequence context of the variants, except the allele change, we extracted the 5-mer flanking sequence centered on the target variant. Reference genomes for the two species in FASTA format were obtained from UCSC (Nassar et al., 2023) and the Samtools faidx command was used to extract the flanking sequence from the reference genome (Li et al., 2009). To calculate the distances from variants to different motifs, scanMotifGenomeWide.pl in HOMER (Duttke et al., 2024) was used to predict motif sites across both the human and cattle genomes. The 460 known vertebrate motifs provided by HOMER were used as input for genome-wide scanning. After obtaining the genome-wide positions of the motifs, bedtools closest (Quinlan & Hall, 2010) was used to calculate distances.

In addition to general annotations, 5313 predicted functional genomic data for each variant were obtained using the Enformer model (Avsec et al., 2021). This includes a diverse set of functional genomic signals, including chromatin accessibility (ATAC-seq, DNase-seq), histone modifications (ChIP-seq), transcription factor binding sites, and gene expression (CAGE-seq). The pre-trained Enformer model was loaded from Tensorflow hub (https://tfhub.dev/deepmind/enformer/1), and the reference genome in FASTA format for the target species was downloaded from UCSC (Nassar et al., 2023). Python code for running the pre-trained Enformer model was adopted from the Enformer GitHub page (https://github.com/deepmind/deepmind-research/tree/master/enformer). The pre-trained model was employed to make predictions for the sequences (393,216 bp) centered on the reference and alternative alleles, respectively, where 393,216 bp represents the sequence window used as input to Enformer. Subsequently, the 5313-column scores for each variant were defined as the differences between the reference and alternative prediction results.

To facilitate the reusability of the aforementioned annotation approaches, a SNV annotation pipeline was developed using the Nextflow framework (Di Tommaso et al., 2017). These annotation approaches were encapsulated into separate processes, which can run in parallel to enhance annotation efficiency. They were organized into two sub-workflows, one for general annotations and another for Enformer scores. The default values of parameters and profiles for various execution environments were defined in the configuration files. The pipeline can be accessed from: https://github.com/evotools/nf-VarAnno.

### Genotype datasets

Various variant sets were used in the study to explore the utility of annotations in regulatory variant modelling. Human fine-mapped regulatory variants were obtained from the CaVEMaN dataset from the Genotype-Tissue Expression (GTEx) project (GTEx Consortium, 2020). We focused on autosomal SNPs, resulting in a set of 590,778 regulatory variants (foreground data) for analysis. For obtaining high-confidence regulatory variants, we filtered the 590,778 variants to those with a CaVEMaN causal probability > 0.5, resulting in 45,987 variants, where CaVEMaN is a statistical framework designed to prioritize causal variants (Brown et al., 2017). To maintain consistency with the regulatory variants, matching sets of background variants were down-sampled from variants located within 1Mb of the TSS and selected to match the minor allele frequency (MAF) distribution of the regulatory variants. The MAF of human variants were obtained from VEP (McLaren et al., 2016a). Additionally, to benchmark the performance of our approach, we obtained 11.4k variants, which include 1,140 putative functional regulatory variants across 83 human complex traits and 10,260 control variants, used in a recent paper benchmarking models for causal regulatory variant prediction in human (Benegas, Eraslan, et al., 2025).

Similarly, regulatory variants from cattle GTEx were acquired for cattle modelling (Liu et al., 2022). Lead variants, denoting those with the smallest nominal p-value for each gene across all tissues, were extracted. Furthermore, these variants were filtered based on their associated permutation p-values, using a threshold of 0.05. After filtering, 79,215 SNPs were retained as regulatory. Matching sets of background variants were down-sampled from the remaining dataset, selecting only variants with a nominal p-value greater than 0.05 and ensuring that their MAF distribution matched that of the foreground variants. Cattle MAF was calculated using BCFtools based on the reference panel from cattle GTEx (Liu et al., 2022).

For both humans and cattle, we obtained another set of regulatory variants (raQTL) based on the survey of regulatory elements (SuRE) technique. The human raQTLs were obtained from van Arensbergen et.al. (van Arensbergen et al., 2019). The original variants on hg19 were lifted over to hg38, resulting in 18,717 SNPs in the K562 cell line and 13,838 SNPs in the HepG2 cell line. The same number of background variants for these two cell lines were randomly sampled from the tested variants with SuRE signals greater than 4 and p-values greater than the threshold for defining raQTLs in the original paper (0.006192715 for K562 and 0.00173121 for HepG2). The cattle raQTLs were generated following the same approach as in van Arensbergen et.al. and is described in full detail in an accompanying paper. Instead of using the statistical approaches employed for identifying raQTLs in humans, our group used zero-inflated negative binomial regression models to analyze SuRE read count files. After filtering according to the adjusted p-value with a threshold of 0.05, there were 66,501 variants kept for further analysis. The same number of matching background variants were randomly sampled from the tested variants with adjusted p-value greater than 0.05.

To assess the effectiveness of human and cattle GTEx models in predicting regulatory effects of reported variants associated with traits, we used data from genome-wide association studies. For human GWAS data, we used the variants from the credible sets in Open Target Genetics (Ghoussaini et al., 2021). We downloaded the datasets from Open Target Genetics and the Python package pyarrow (Richardson et al., 2023) was employed to convert the data into a Python dataframe from parquet format. Afterward, the variants were filtered based on a posterior probability threshold of 0.5, and those variants also present in the human high-confidence training set were removed, resulting in 28,918 variants. For comparison purposes, an equal number of background variants were randomly selected from the remaining variants with posterior probability smaller than 0.5. Another set of pathogenic regulatory variants associated with Mendelian diseases was obtained from Damina et al. (Smedley et al., 2016). The 406 single nucleotide variants were initially lifted over from the GRCh37 assembly to the GRCh38 assembly using the UCSC liftOver tool (Kuhn et al., 2013). After excluding those on sex chromosomes, 345 SNVs were retained for analysis. An equal number of background variants were randomly selected from the human 1000 genomes cohort (1000 Genomes Project Consortium et al., 2015).

For cattle GWAS data, we initially obtained the GWAS summary statistics data for three cattle traits: body condition score, body weight, and stature, from Edwardo et al (Reynolds et al., 2021). The originally reported variants on cattle genome UMD3.1 were lifted over to ARS-UCD1.2 using the mapping file provided in the paper, successfully mapping 98.6% of the variants between the assemblies. Subsequently we conducted conditional analysis using GCTA-COJO to select independently associated SNPs for each trait (Yang et al., 2012). To run GCTA-COJO, the initial summary statistics data were re-organized to match the input format required by the command. PLINK 2.0 (Chang et al., 2015) was used to generate the PLINK binary PED files, which served as an input for GCTA-COJO. To create the reference VCF files needed for input into PLINK 2.0, we first downloaded the reference VCF files from the 1000 Bull Genomes Project (Hayes & Daetwyler, 2019). Then these files were filtered using bcftools (Danecek et al., 2021) to include only a subset of samples that match the breeds included in the cattle GWAS data. After obtaining the associated SNPs for these traits, they were filtered based on their p-values using a threshold of 5e-8. Consequently, 96, 344 and 241 associated variants were retained for body condition score, body weight, and stature, respectively. Additionally, we obtained fine-mapping results for milk yield from Jiang et al. (Jiang et al., 2019). The variants from the cattle genome UMD3.1 were lifted over to ARS-UCD1.2, the allele changes were adjusted based on the liftOver chain file, and a p-value threshold of 5×10^-8^ was applied to identify 210 associated variants for milk yield. For comparison, we obtained background data with p-values greater than 0.05, selecting the same number of variants as the foreground 100 times for each trait.

### Regulatory variant modelling

The variants were annotated using the variant annotation pipeline. These annotations first underwent feature engineering processes, performed using the Python scikit-learn package (Pedregosa et al., 2011). The conservation score annotations contained missing values, affecting approximately 0.9% of human variants and 0.8% of cattle variants. Given the small proportion, we opted to exclude variants with missing conservation scores from the dataset rather than employing imputation strategies. Subsequently, categorical features were encoded using different strategies. For sequence categorical features, such as 5-mer flanking sequence and allele change, we created a dictionary to map each nucleotide in the sequence to a binary string (A: 1000, C: 0100, G: 0010, T: 0001). The resulting string was then divided into separate integer columns, each indicating the presence or absence of a particular base. Categorical features without an ordinal relationship, such as chromosome and variant consequence, were encoded using one-hot encoding approach. After encoding, the transformed features were reintegrated into the feature table, replacing the original categorical features. This resulted in a 1680-column encoded general feature table for human variants and 96-column for cattle variant.

Four machine learning algorithms were considered for model construction, Random Forest (Breiman, 2001), CatBoost (Prokhorenkova et al., 2017), XGBoost (T. Chen & Guestrin, 2016), and Support Vector Machines (SVM) (Cortes & Vapnik, 1995). Random Forest and SVM models were implemented using scikit-learn libraries, while CatBoost and XGBoost utilized their respective Python packages. Considering both training efficiency and model performance, CatBoost was selected as the algorithm for modelling in the study. Each variant set, including both foreground and background variants, was split into training and test sets by chromosomes to prevent data leakage. Specifically, several chromosomes were left out for testing, while the remaining chromosomes were used for model training and hyper-parameter tunning. To optimize hyper-parameters, we used BayesSearchCV from the Python scikit-optimize library (Head et al., 2018), a Bayesian optimization-based approach that efficiently tunes hyper-parameters through iterative cross-validation. For cross-validation, we implemented chromosome-level cross-validation, where entire chromosomes were held out in each fold to ensure that variants from the same chromosome did not appear in both training and validation sets. When performing benchmarking, we used a single-chromosome hold-out strategy for hyper-parameter tuning and final evaluation. An incremental learning strategy was adopted to leverage human data for cattle variant prediction (Wu et al., 2019). The human model, trained on human annotations available for both human and cattle datasets, was employed as the initial model for a subsequent training step utilizing cattle annotations. For the comparison of models with and without Enformer features, we trained these models using the CatBoost algorithm with default hyperparameters due to limitations in available computational resources. The model tuning and training were conducted using GPUs from the Edinburgh international data facility (EIDF).

To evaluate the models’ performance beyond just accuracy, various metrics were calculated using scikit-learn packages, including precision, recall, F1-score, and the Area Under the Curve (AUROC), to assess their proficiency in true positive predictions and discriminative capacity between regulatory variants and other variants. Furthermore, the Shapley Additive exPlanations (SHAP) approach was employed to elucidate the relationship between feature values and model outputs, thereby enhancing model interpretation (S. Lundberg & Lee, 2017).

## Code availability

The cross-species variant annotation pipeline is available on GitHub: https://github.com/evotools/nf-VarAnno. The machine learning pipeline is available on GitHub: https://github.com/evotools/functional-variant-prediction.

## Data availability

The cattle conservation metrics are available at: https://doi.org/10.5281/zenodo.13332541. The cattle tissue-specific and cross-tissue models for functional variant prediction can be found at: https://doi.org/10.5281/zenodo.14901001.

## Acknowledgments

This work was supported by grant BB/W000288/1 and Institute Strategic Programme Grant BBS/E/D/10002070 from the Biotechnology and Biological Sciences Research Council (BBSRC). This project was also supported by funding by the Bill and Melinda Gates Foundation (BMGF) and with UK aid from the UK Government’s Department for International Development (Grant Agreement OPP1127286) under the auspices of the Centre for Tropical Livestock Genetics and Health (CTLGH), established jointly by the University of Edinburgh, SRUC (Scotland’s Rural College), and the International Livestock Research Institute (ILRI). The findings and conclusions are those of the authors and do not necessarily reflect the positions or policies of the BMGF or the UK Government. We would like to thank Dr Ivan Pocrnic for helpful discussions regarding this work.

## Supplementary materials

**Supplementary Figure 1.**
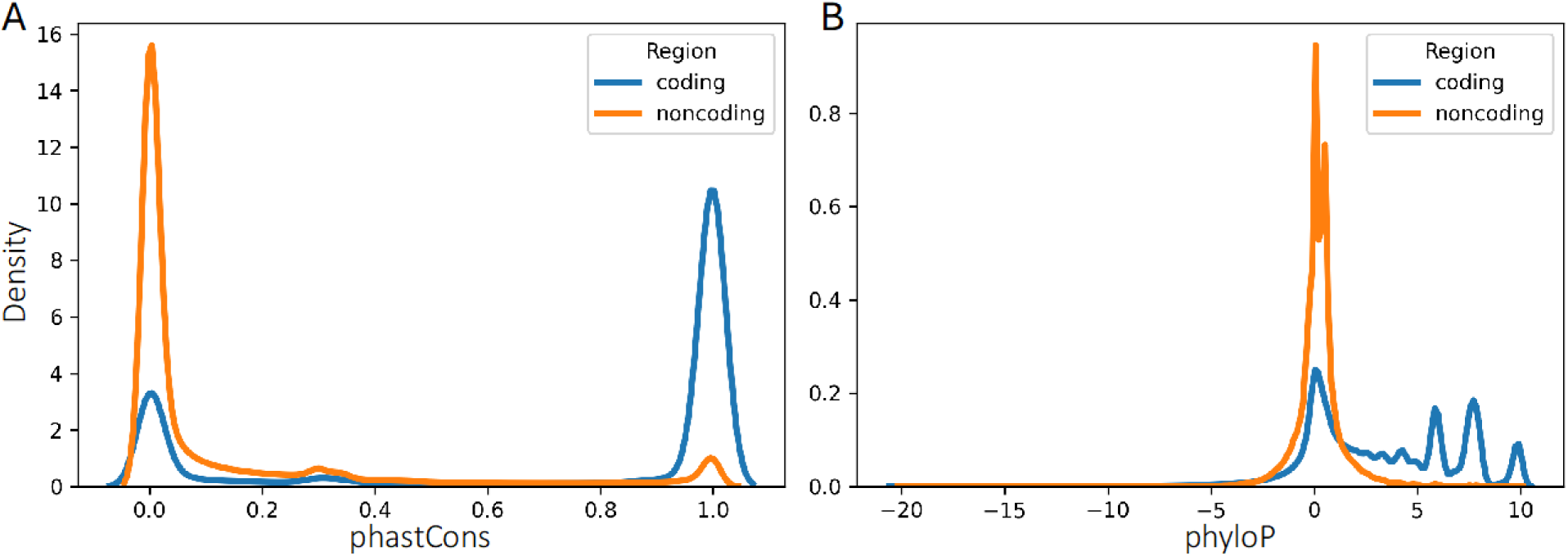
Conservation score distribution differences in coding and noncoding regions in the cattle genome. (A) phastCons241way conservation score (B) phyloP241way conservation score.

**Supplementary Figure 2.**
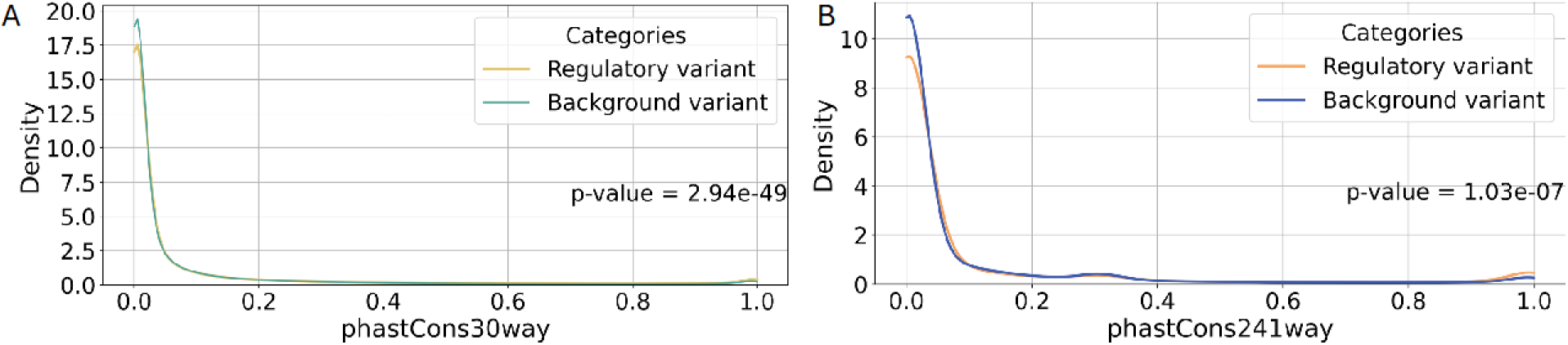
Kernel density estimate (KDE) of the distribution of phastCons scores for (A) human and (B) cattle. The conservation scores display significant differences between the groups in both human and cattle (Two-sample Kolmogorov-Smirnov test p-values are shown in the corresponding plots).

**Supplementary Figure 3.**
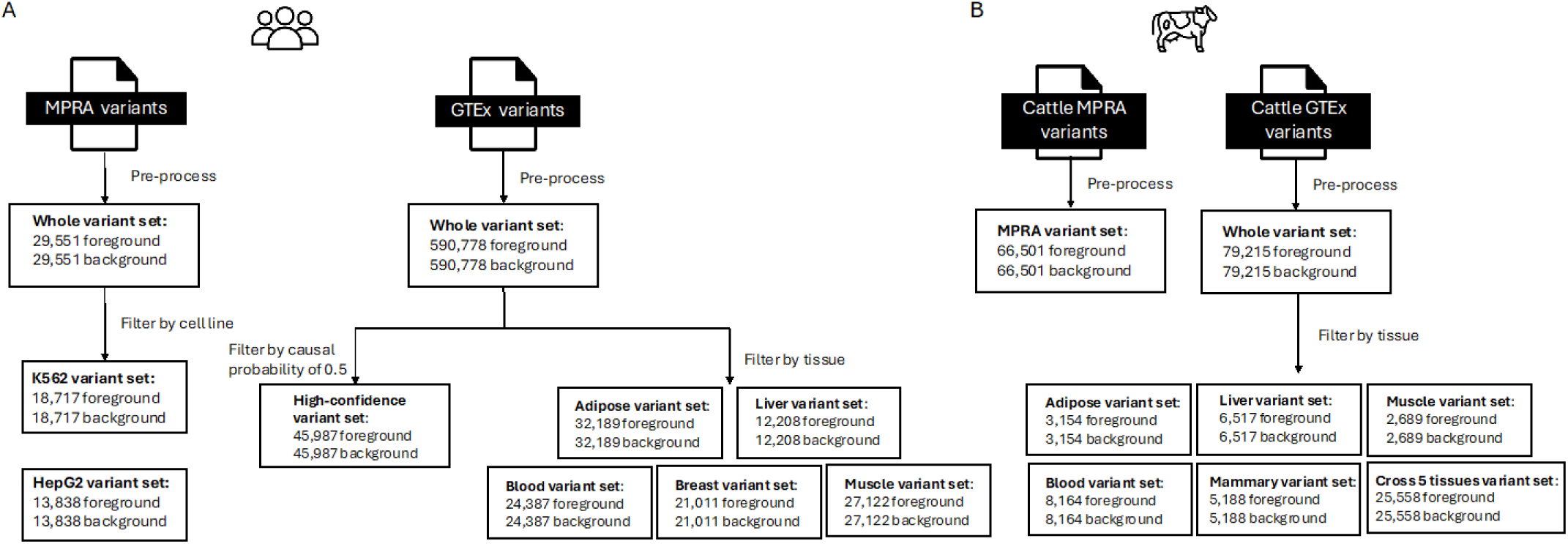
An overview of different variant sets included in the study. (A) Human. (B) Cattle

**Supplementary Figure 4.**
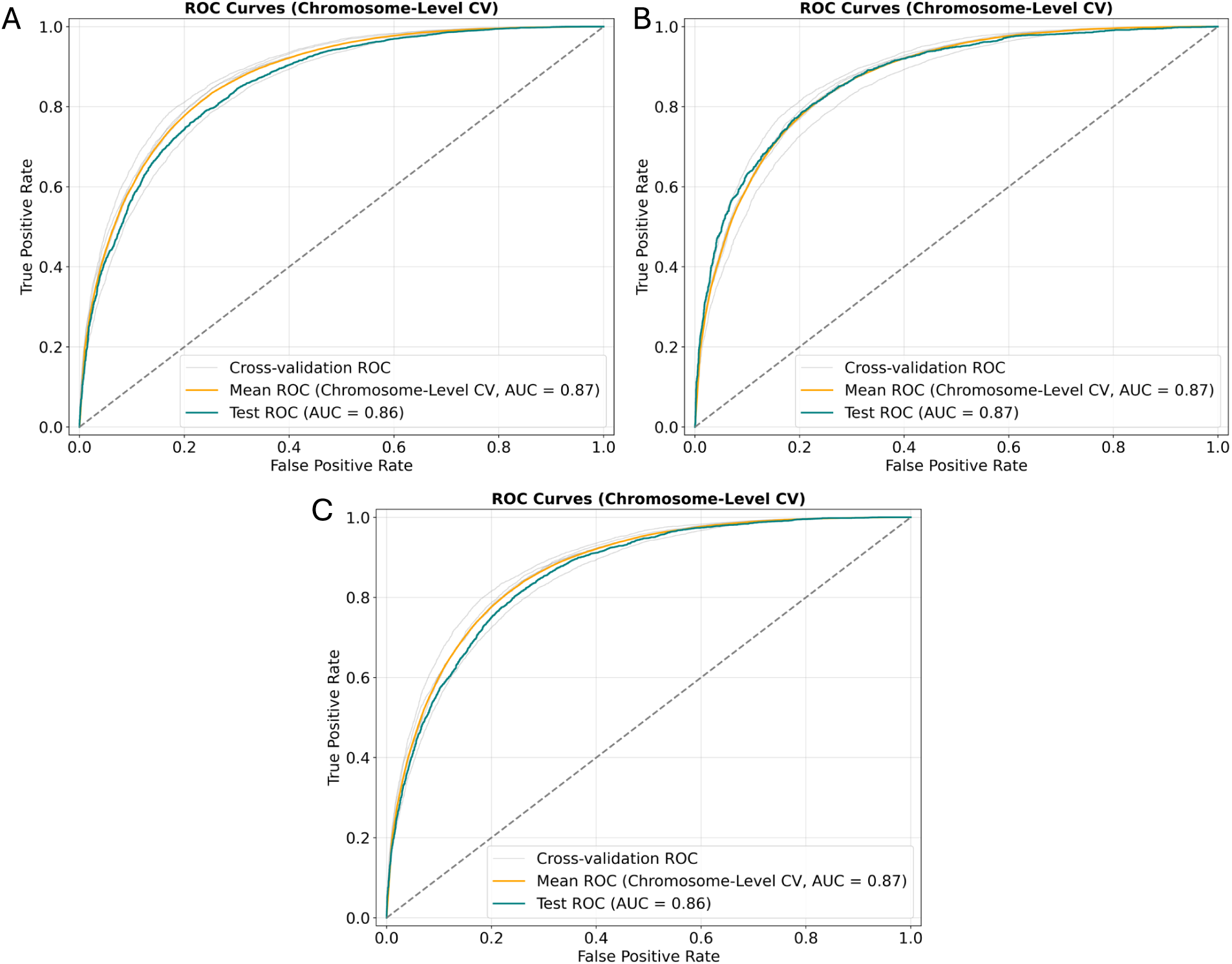
ROC curves and AUROC (AUC) scores for the human high-confidence model when testing on (A) chr1 and chr22, (B) testing on chr3, and (C) testing on chr2 and chr21.

**Supplementary Figure 5.**
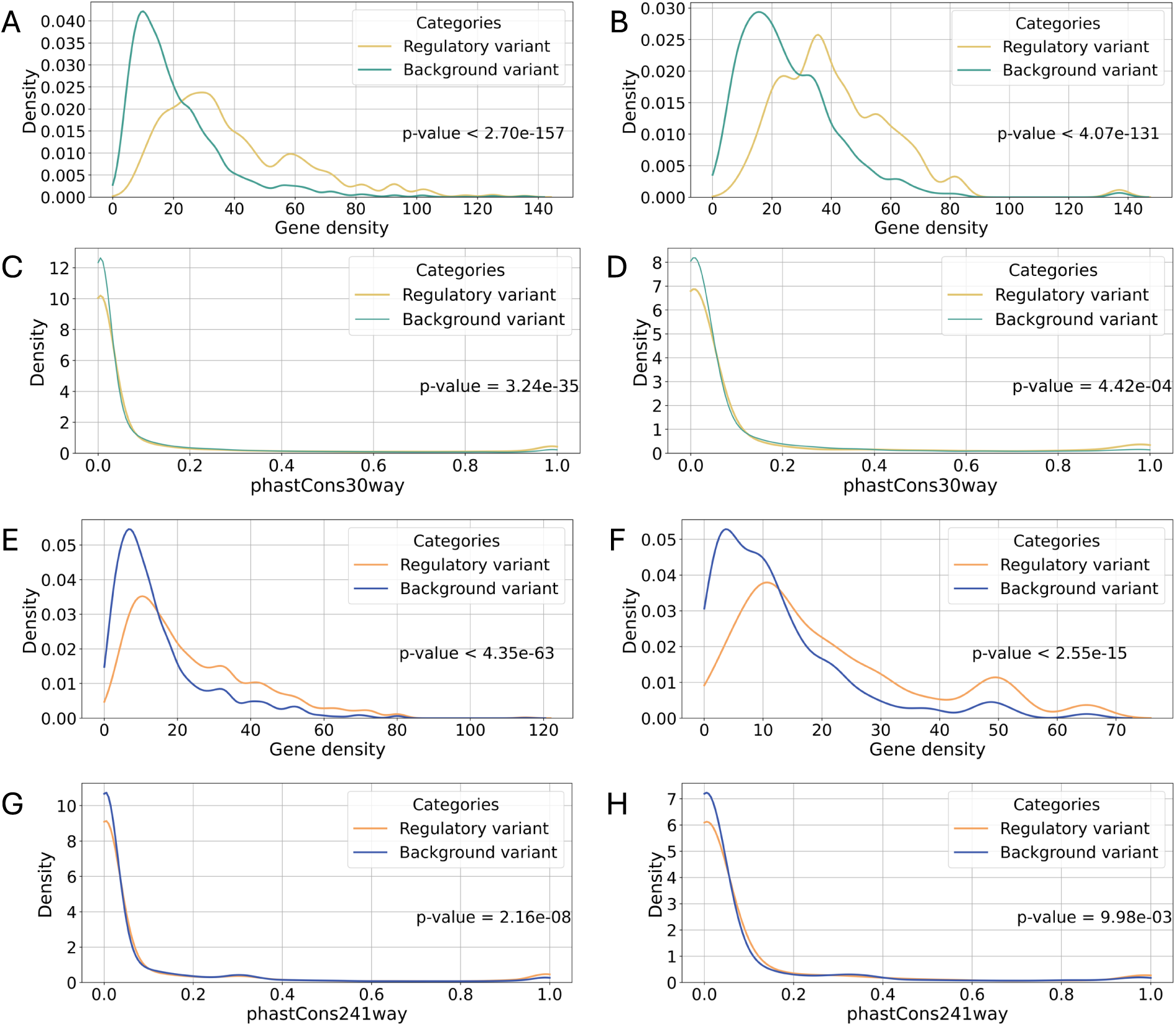
Feature distributions of foreground and background variants in training and test sets. (A) Human gene density distribution in the training set. (B) Human gene density distribution in the test set. (C) Human phastCons30way distribution in the training set. (D) Human phastCons30way distribution in the test set. (E) Cattle gene density distribution in the training set. (F) Cattle gene density distribution in the test set. (G) Cattle phastCons241way distribution in the training set. (H) Cattle phastCons241way distribution in the test set. All distributions display significant differences between the groups in both human and cattle (Two-sample Kolmogorov-Smirnov test p-values are shown in the corresponding plots).

**Supplementary Figure 6.**
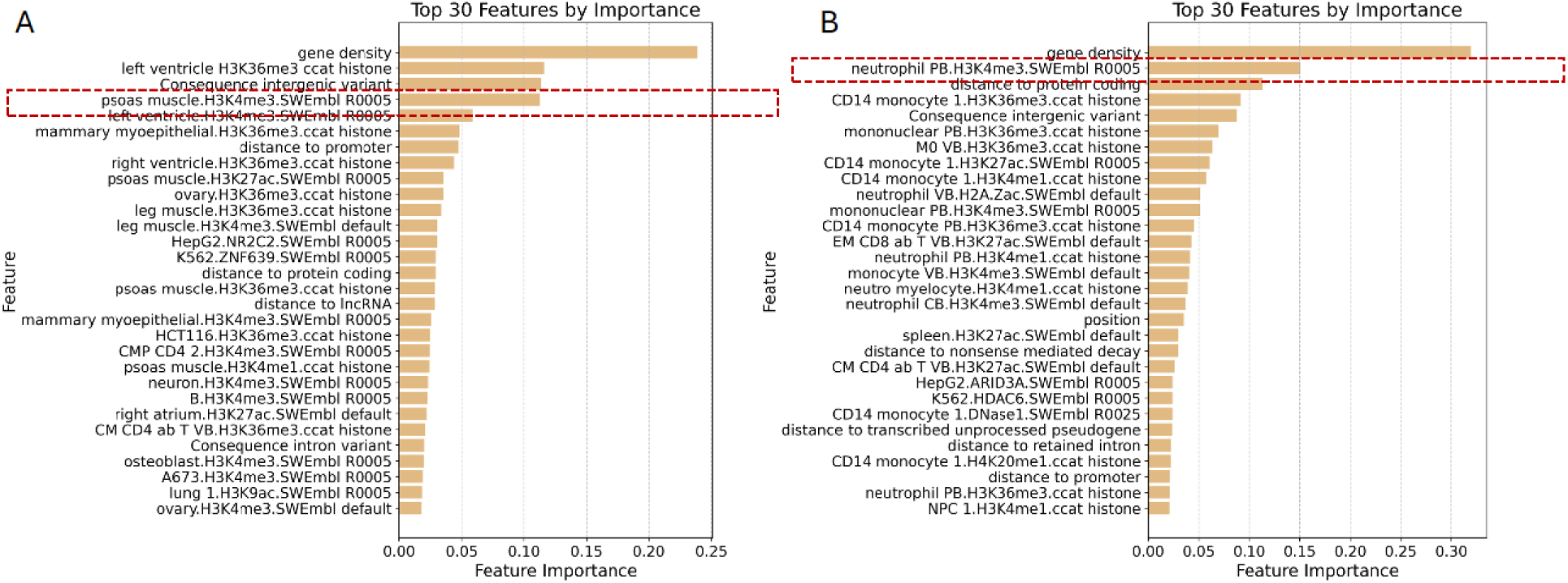
Top 30 features in human (A) muscle and (B) blood specific models. The features are ranked in descending order according to their relative feature importance in the model. The features within the red dashed rectangle are the top chromatin features that exclusively emerged among the top 10 in their respective tissue-specific models.

**Supplementary Figure 7.**
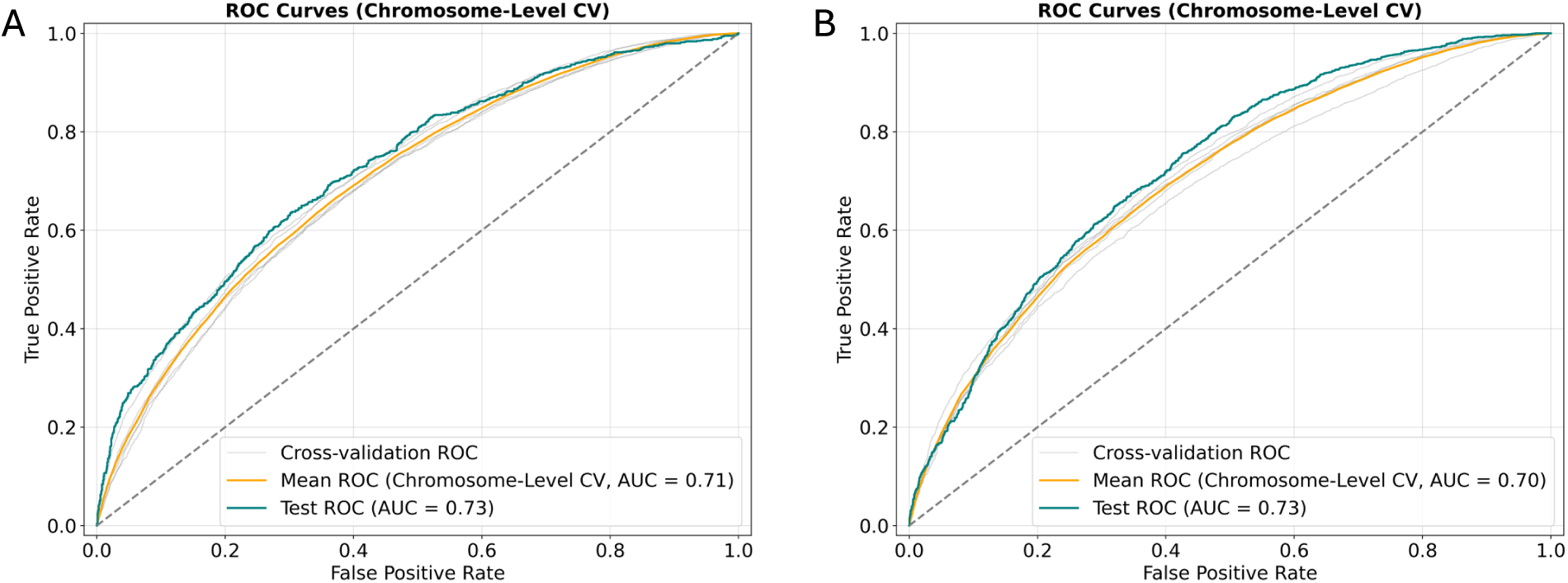
ROC plots and AUROC (AUC) scores for cattle cross-tissues model when testing on (A) chr1 and chr28, (B) testing on chr10.

**Supplementary Figure 8.**
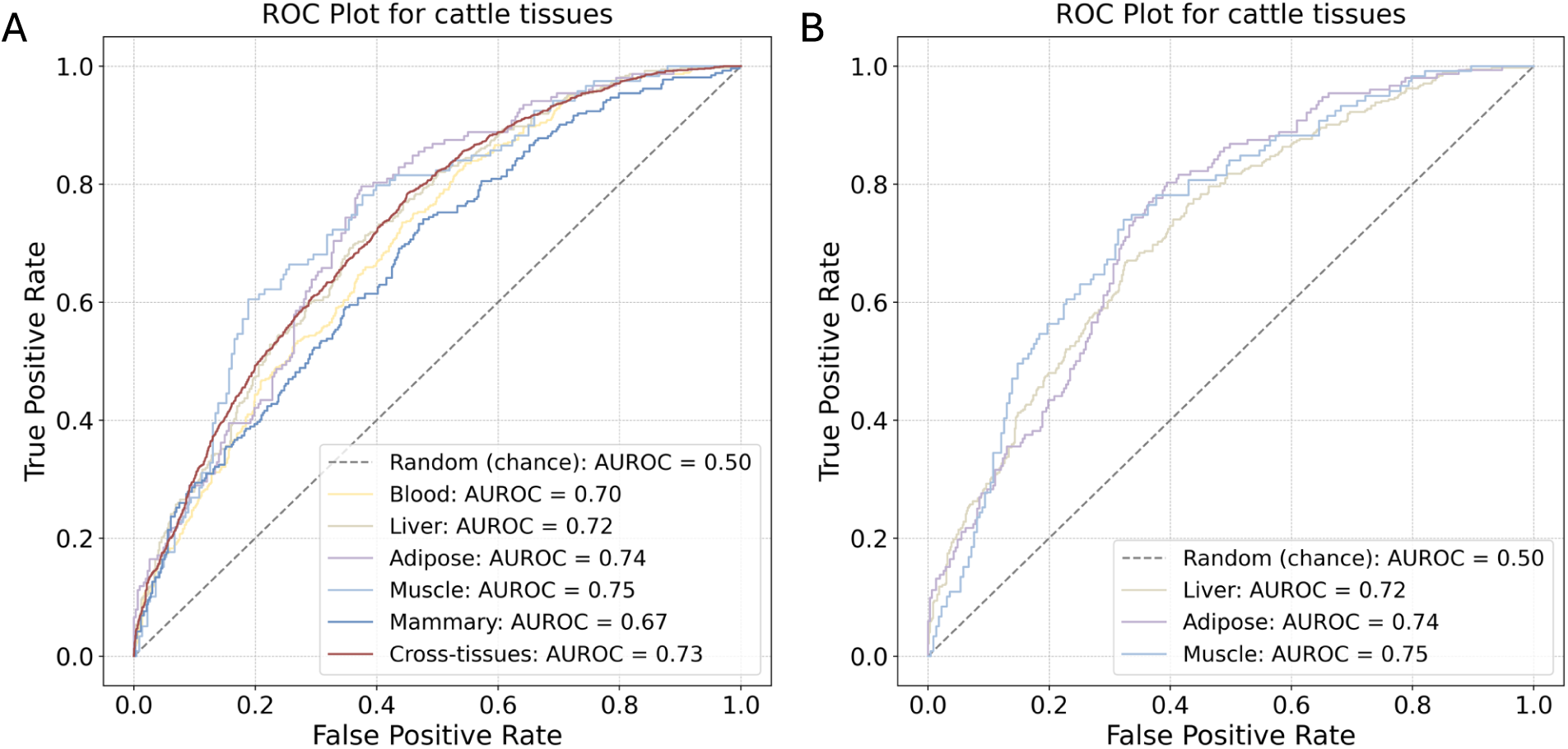
ROC plots and AUROC scores for cattle model trained with (A) full set of 95 cattle chromatin data, and (B) with tissue-specific chromatin data for liver, adipose, and muscle models.

**Supplementary Figure 9.**
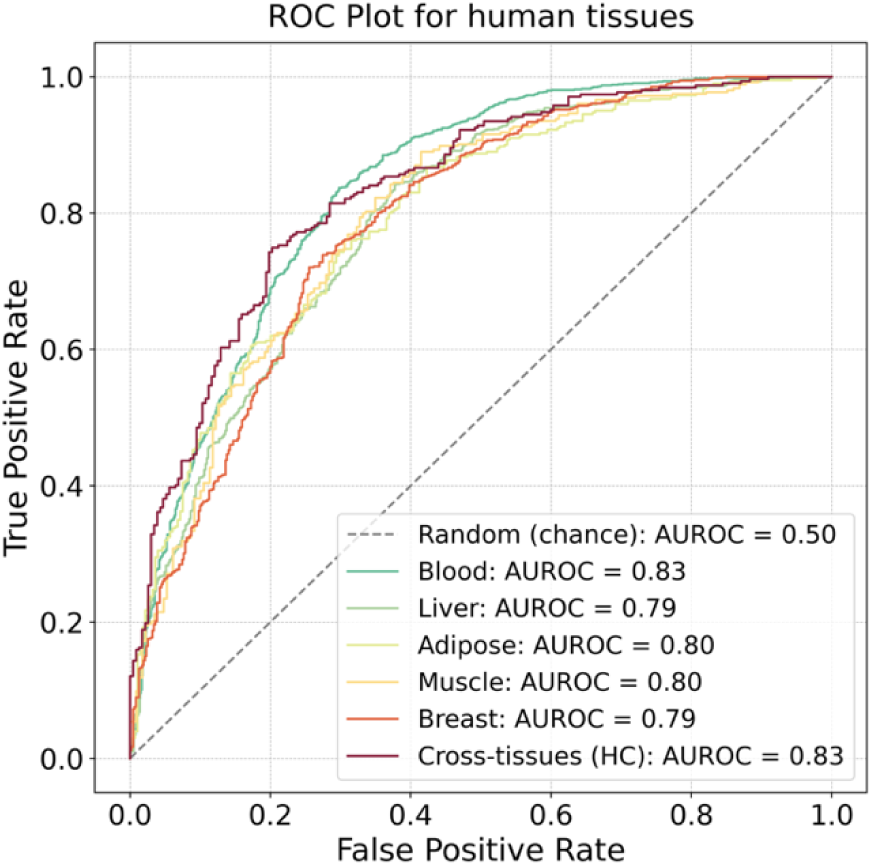
ROC plots and AUROC scores for human models trained and tested with the same number of variants as the corresponding cattle models.

**Supplementary Figure 10.**
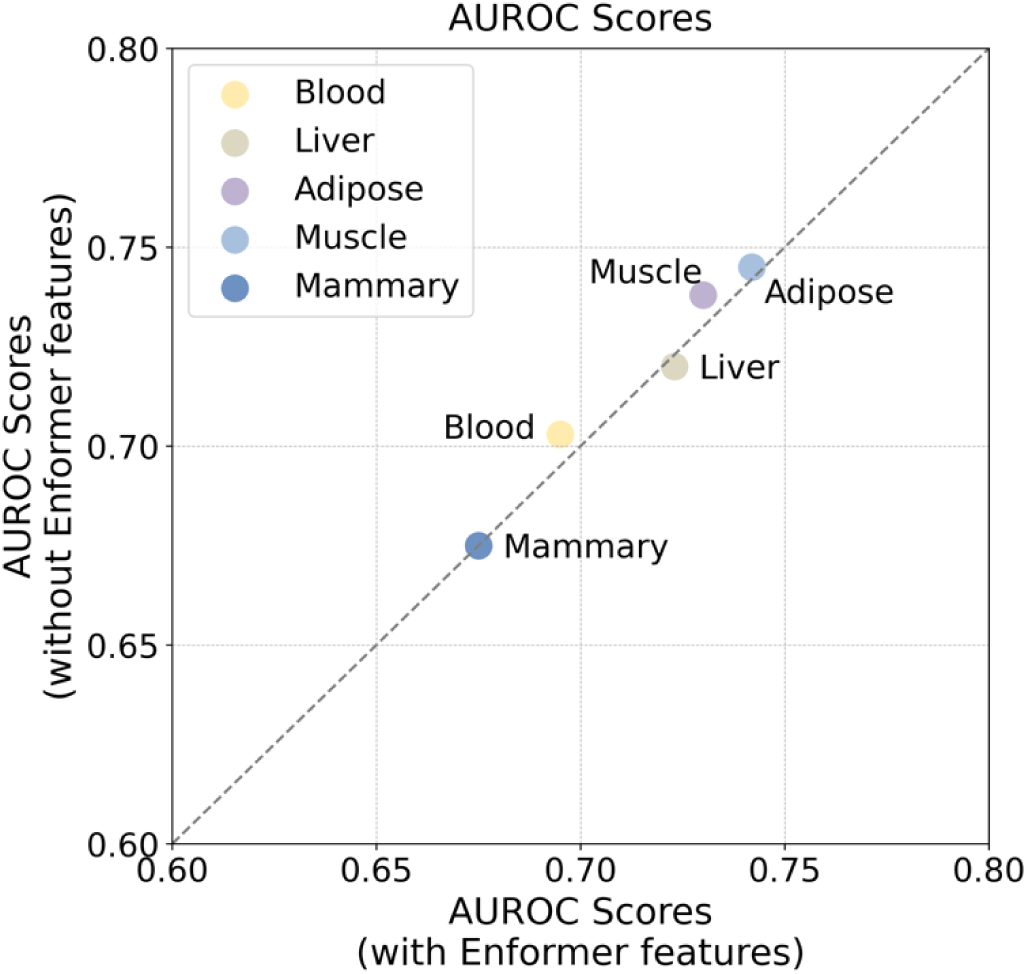
The comparison of the AUROC scores of cattle tissue-specific models trained with or without the full set of Enformer features. The grey dashed line represents parity. The X and Y axis are limited from 0.60 to 0.80 for better clarity of the scatter plot.

**Supplementary Figure 11.**
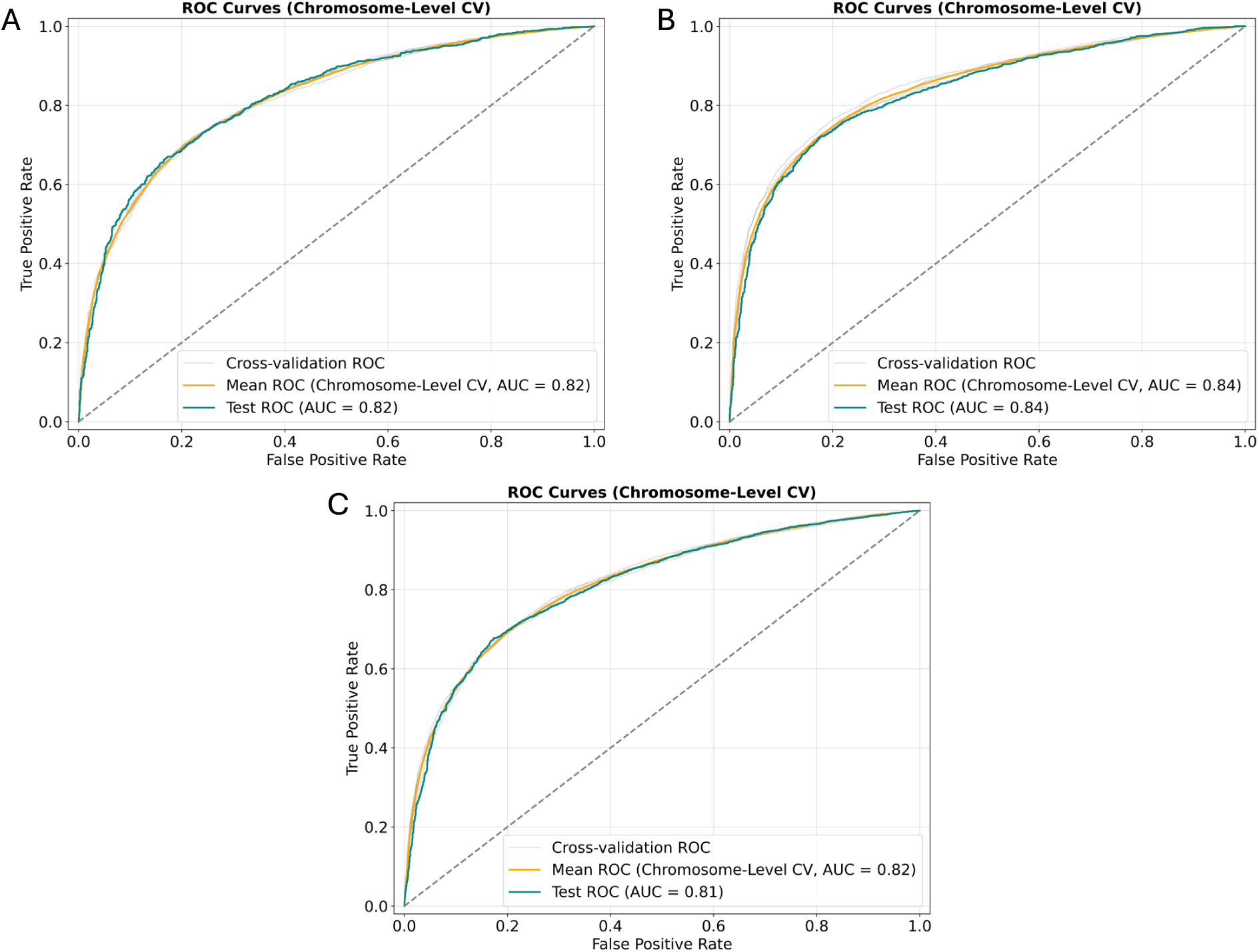
ROC plots and AUROC (AUC) scores for human raQTL models trained and tested in (A) HepG2 cell line. (B) K562 cell line, and (C) both cell lines.

**Supplementary Figure 12.**
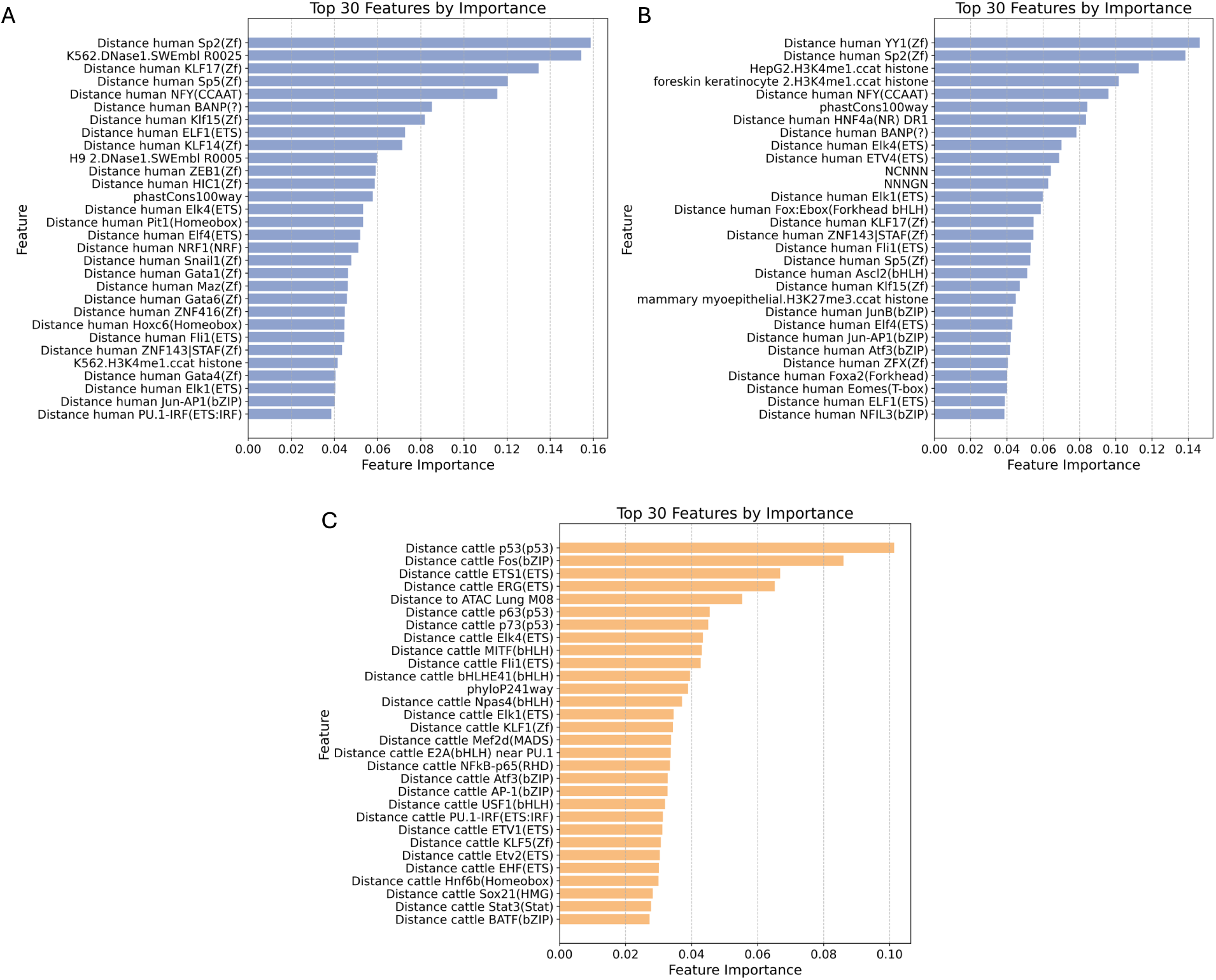
Top 30 features in human and cattle raQTL models (A) K562 cell line, (B) HepG2 cell line and (C) cattle endothelial cell line. The features are ranked in descending order according to their relative feature importance in the model.

**Supplementary Figure 13.**
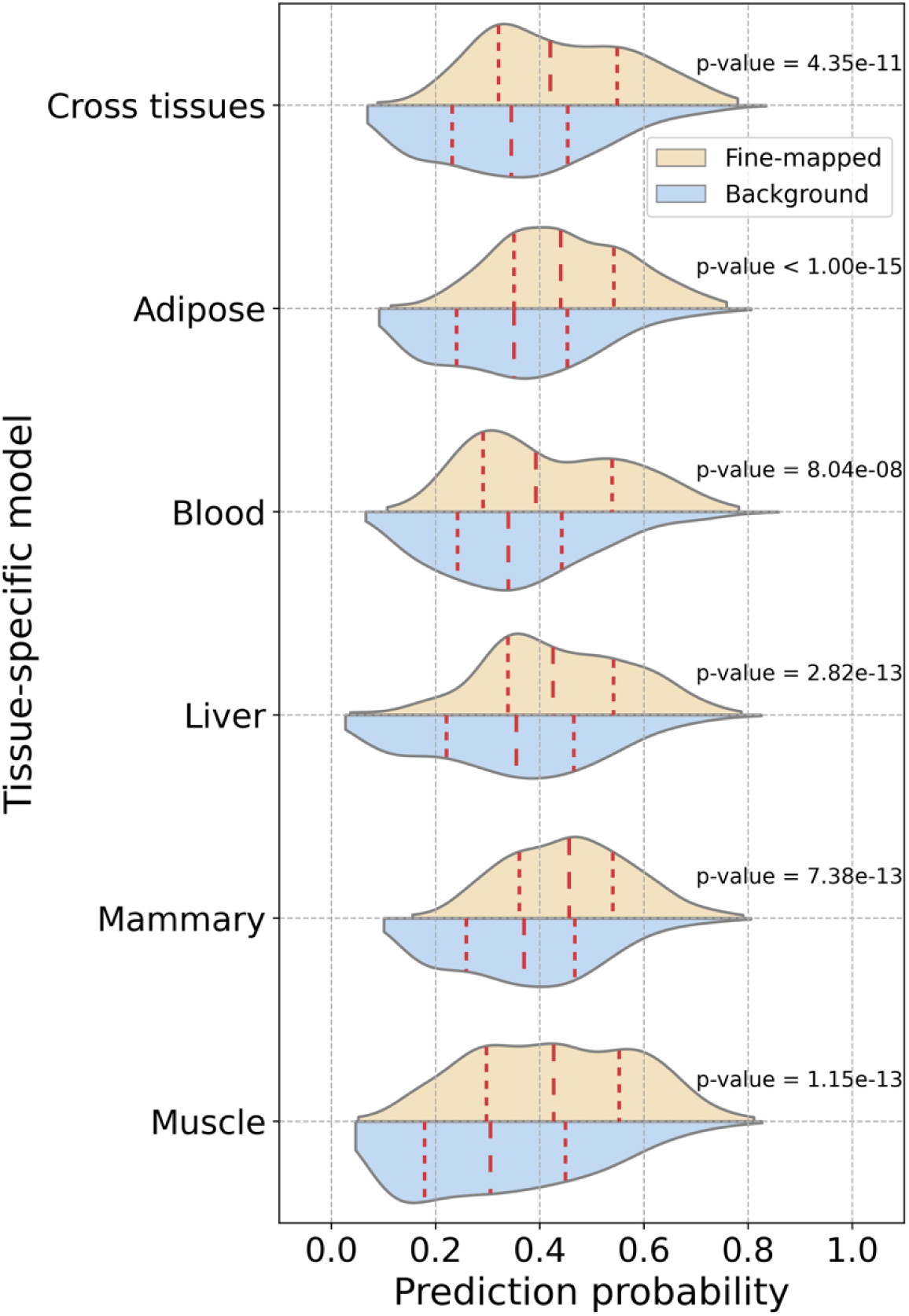
Distributions of model derived prediction probabilities of cattle GWAS fine-mapped and background variants for stature. The red dashed lines in the violin plots represent the median and quartile values. All prediction probabilities are significantly different between fine-mapped and background groups in each experiment. Their corresponding p-values are displayed in the figure (Two-sample Kolmogorov-Smirnov test).

**Supplementary Figure 14.**
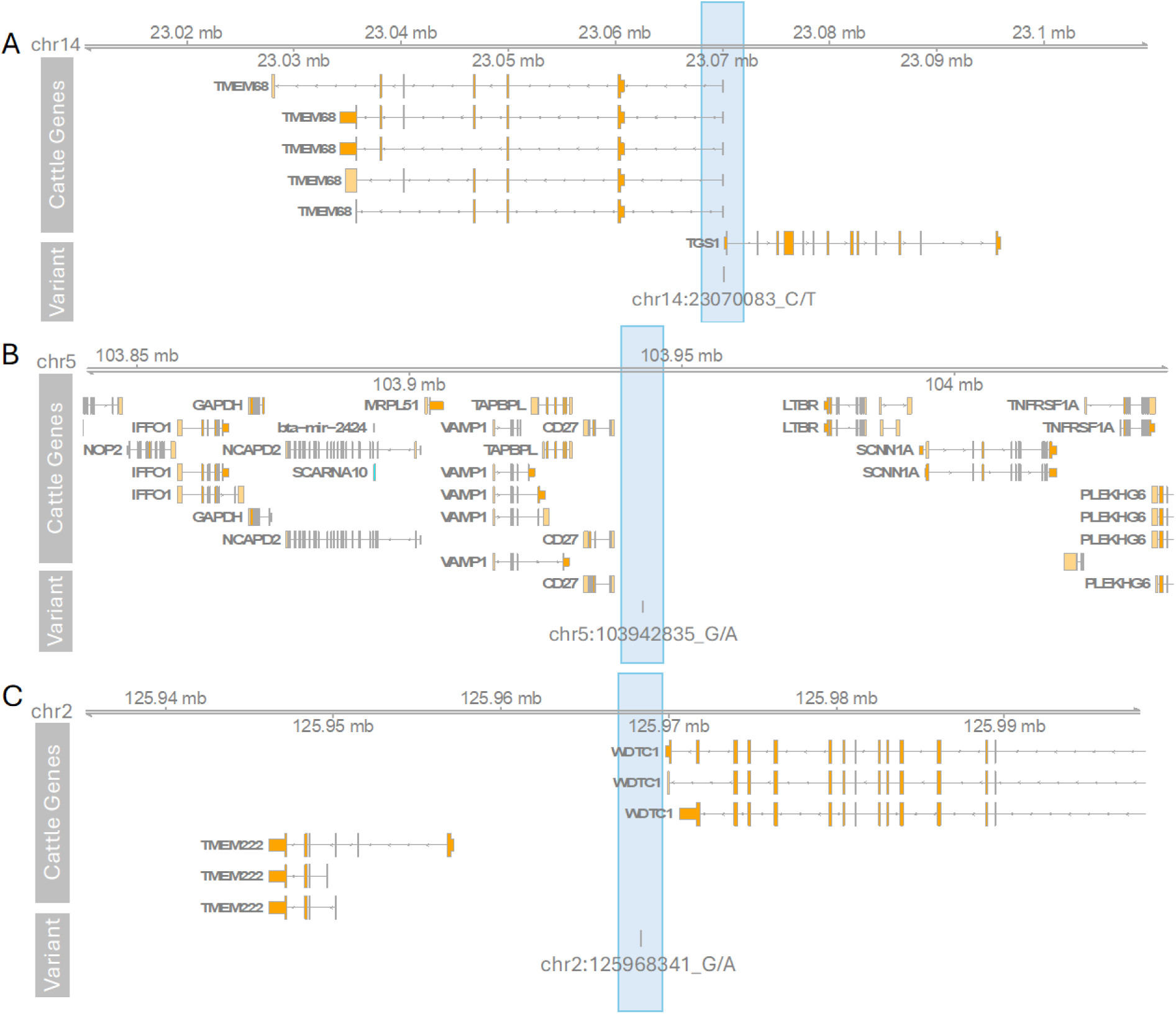
Three cattle trait-associated variants consistently predicted as regulatory variants by all models. (A) stature-associated variant chr14:23070083_C/T, (B) body weight-associated variant chr5:103942835_G/A, and (C) body condition-associated variant chr2:125968341_G/A.

## Notes

### Competing Interest Statement

The authors have declared no competing interest.

### Summary of Updates

In this revised version, we expanded the variant annotation pipeline to include distances to transcription factor motifs (human and cattle) and distances to chromatin data (cattle), resulting in 460 additional human annotations and 555 additional cattle annotations. Incorporation of these features required re-running all analyses, from model training to application. We also modified the train/test split and cross-validation strategy to a leave-out chromosomes design to avoid potential data leakage and adopted a stricter approach to background variant selection. We repeated model training, testing, and trait-associated variant enrichment analyses under this updated framework, including benchmarking the human model against CADD. In addition, we extended the application of our method to a new cattle trait, milk yield, demonstrating its utility in prioritizing variants linked to production phenotypes. Finally, we have made the full machine learning pipeline (feature encoding, model training, and hyper-parameter tuning) available for reuse by the community.

