## Supplementary materials for "The potential of AI models to identify regulatory variants underlying cattle traits"

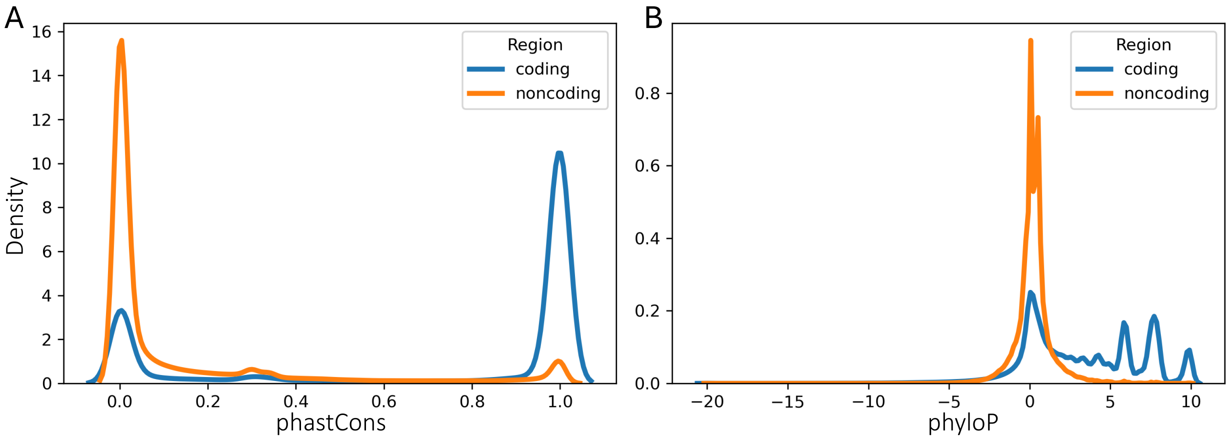


**Supplementary Figure 1 Conservation score distribution differences in coding and noncoding regions in the cattle genome.** (A) phastCons241way conservation score (B) phyloP241way conservation score.

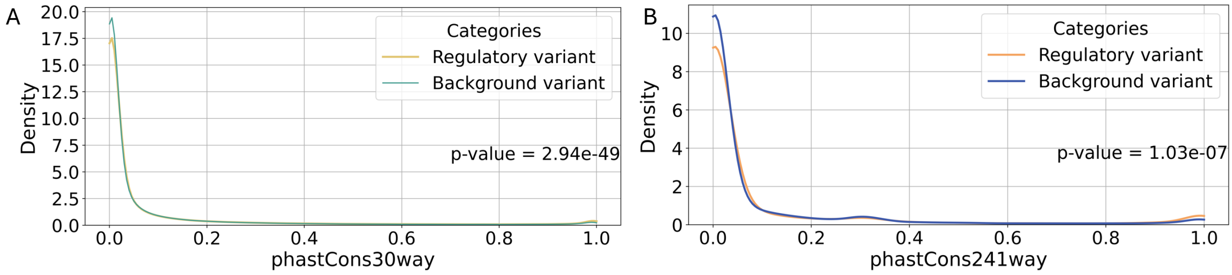


**Supplementary Figure 2 Kernel density estimate (KDE) of the distribution of phastCons scores for (A) human and (B) cattle.** The conservation scores display significant differences between the groups in both human and cattle (Two-sample Kolmogorov-Smirnov test p-values are shown in the corresponding plots).

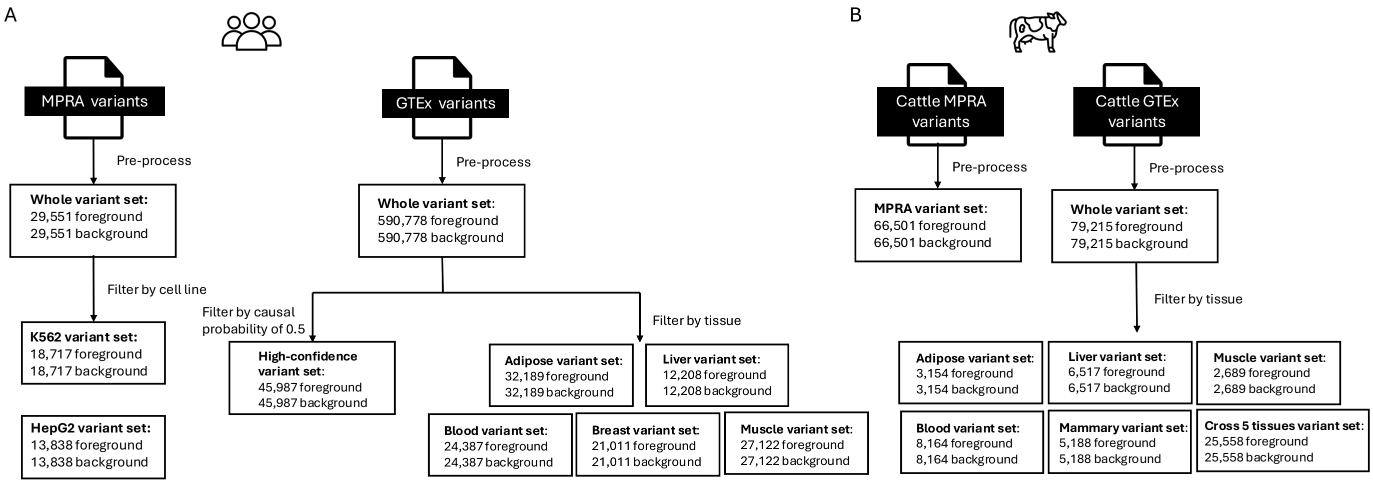

**Supplementary Figure 3 An overview of different variant sets included in the study.** (A) Human. (B) Cattle

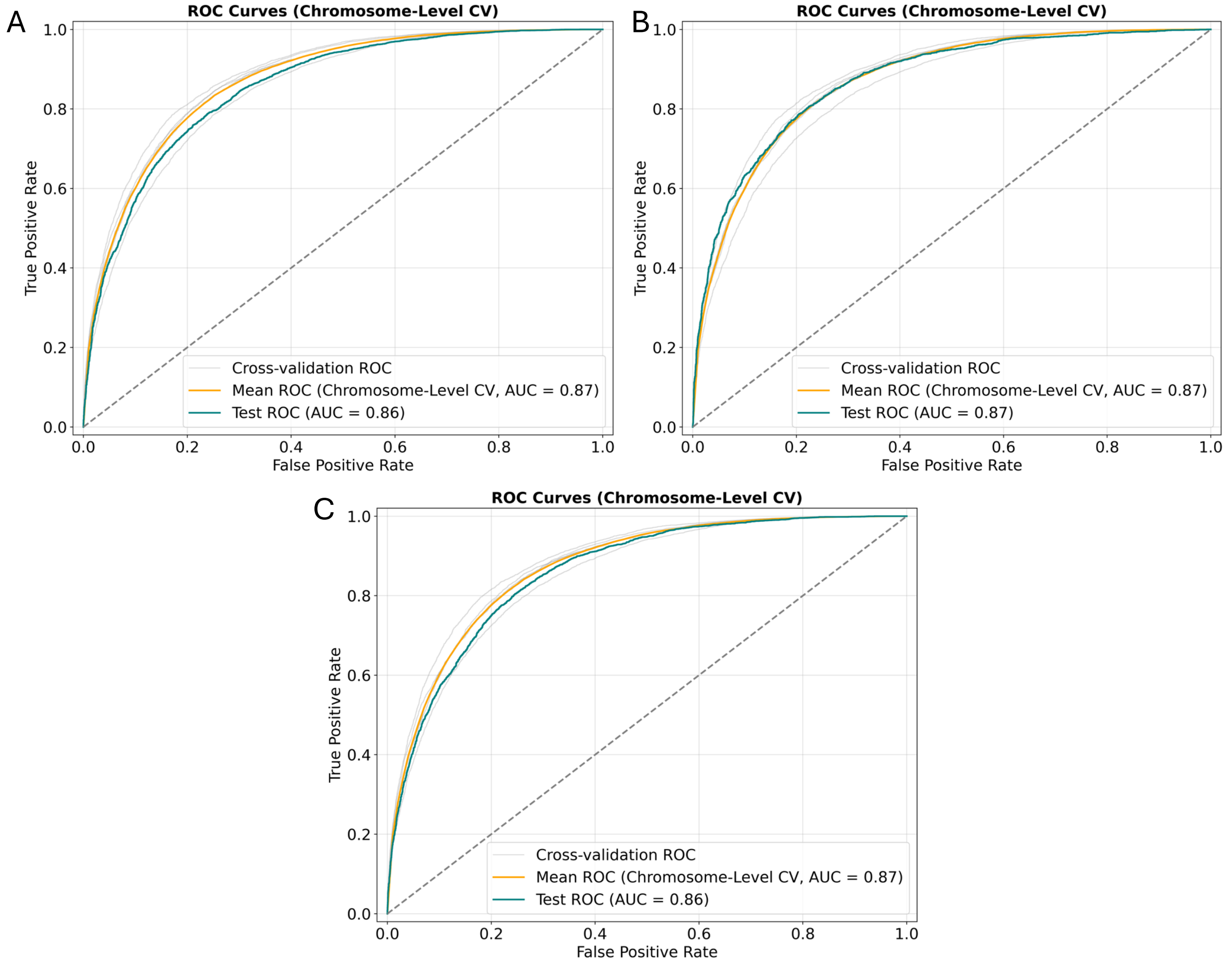


**Supplementary Figure 4** ROC curves and AUROC (AUC) scores for the human high-confidence model when testing on (A) chr1 and chr22, (B) testing on chr3, and (C) testing on chr2 and chr21.

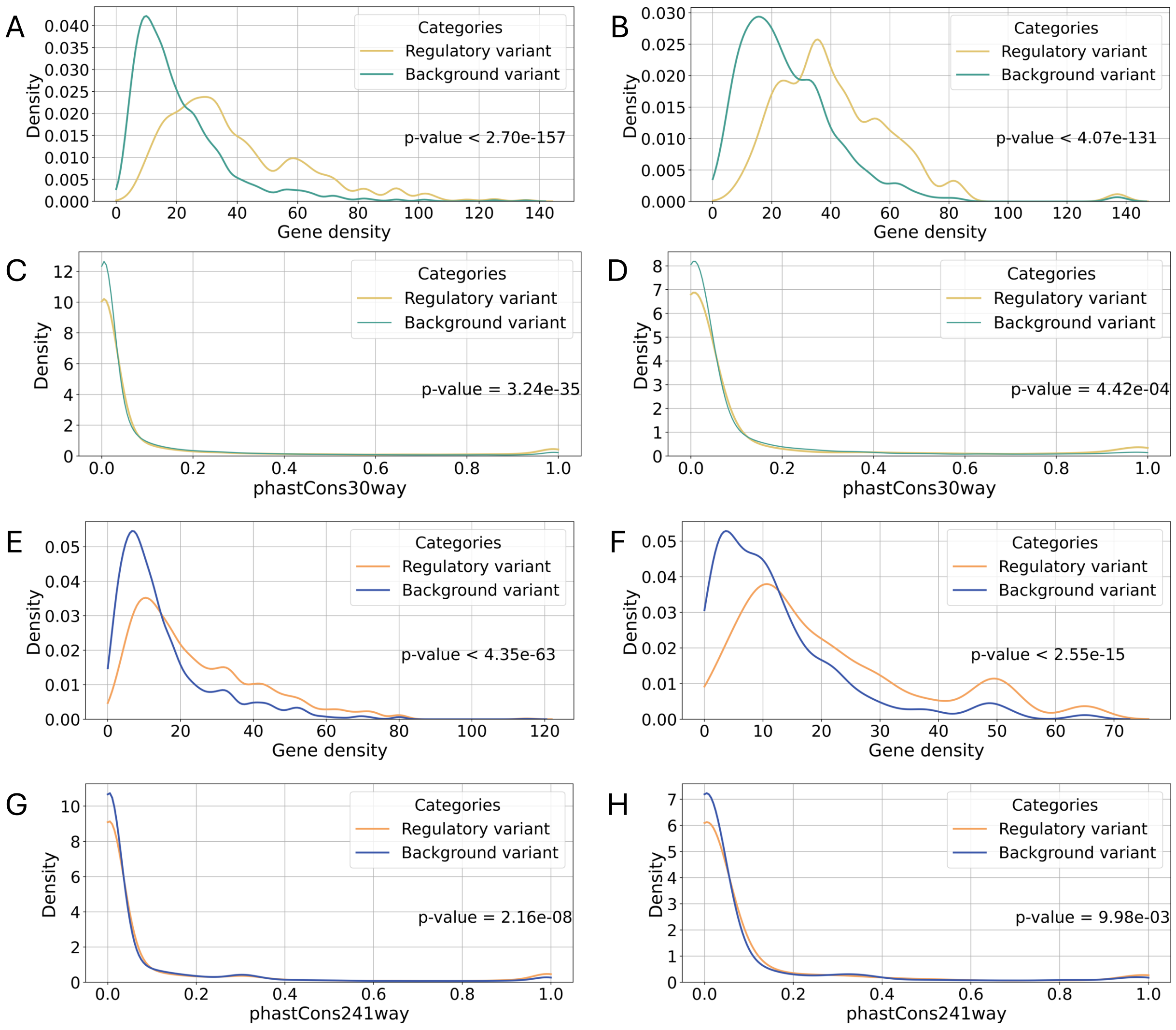


**Supplementary Figure 5 Feature distributions of foreground and background variants in training and test sets.** (A) Human gene density distribution in the training set. (B) Human gene density distribution in the test set. (C) Human phastCons30way distribution in the training set. (D) Human phastCons30way distribution in the test set. (E) Cattle gene density distribution in the training set. (F) Cattle gene density distribution in the test set. (G) Cattle phastCons241way distribution in the training set . (H) Cattle phastCons241way distribution in the test set. All distributions display significant differences between the groups in both human and cattle (Two-sample Kolmogorov-Smirnov test p-values are shown in the corresponding plots).

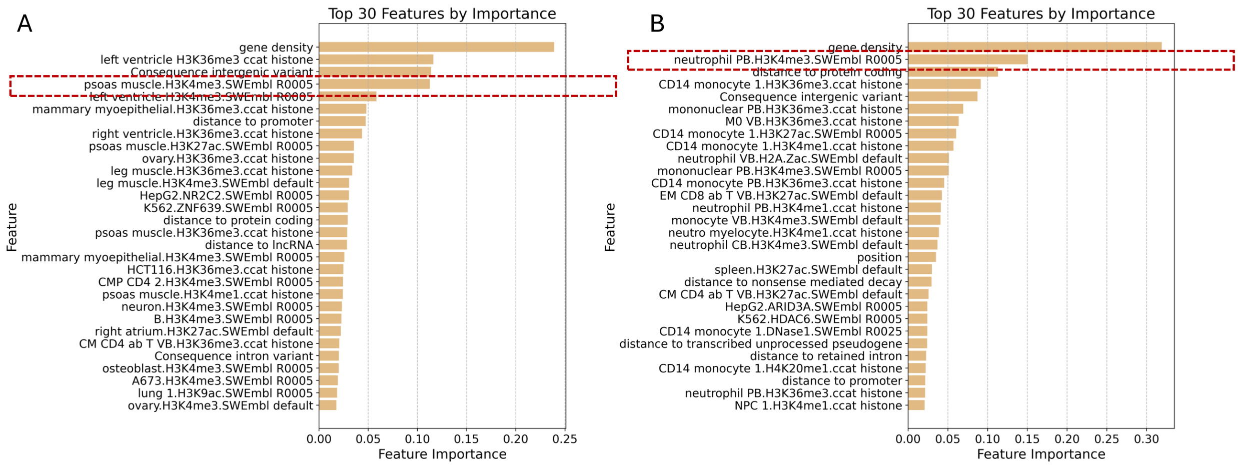


**Supplementary Figure 6 Top 30 features in human (A) muscle and (B) blood specific models.** The features are ranked in descending order according to their relative feature importance in the model. The features within the red dashed rectangle are the top chromatin features that exclusively emerged among the top 10 in their respective tissue-specific models.

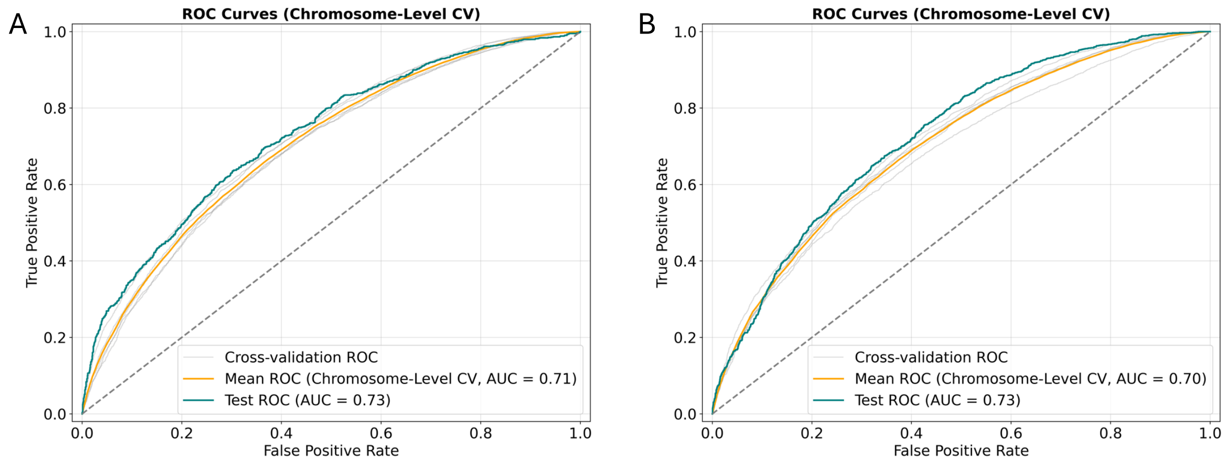


**Supplementary Figure 7** ROC plots and AUROC (AUC) scores for cattle cross-tissues model when testing on (A) chr1 and chr28, (B) testing on chr10.

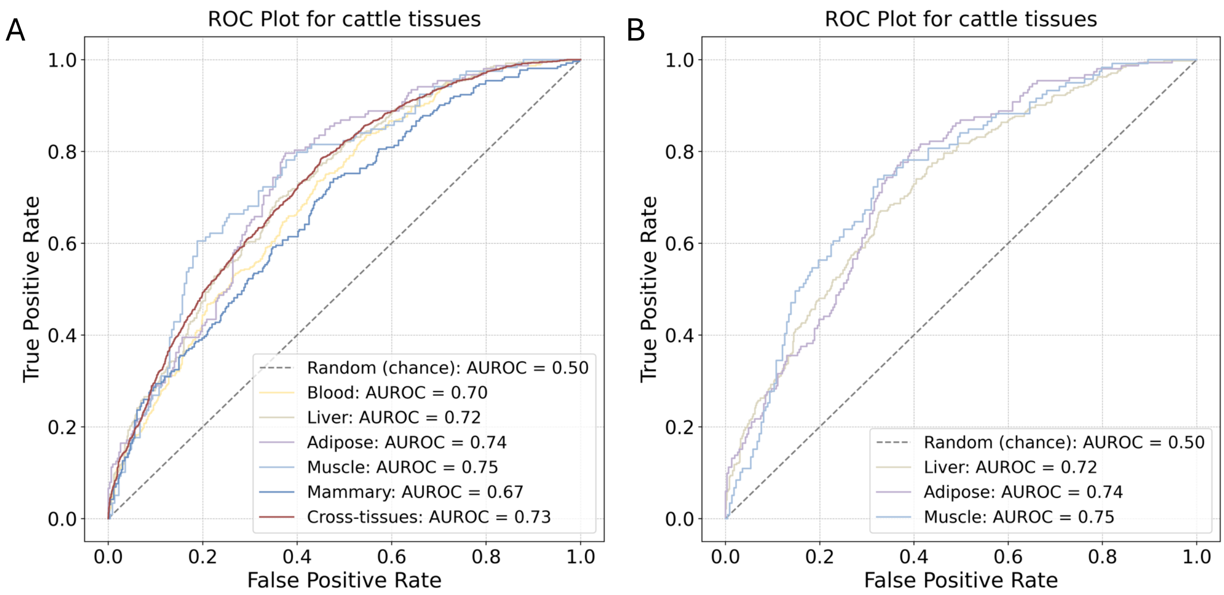


**Supplementary Figure 8** ROC plots and AUROC scores for cattle model trained with (A) full set of 95 cattle chromatin data, and (B) with tissue-specific chromatin data for liver, adipose, and muscle models.

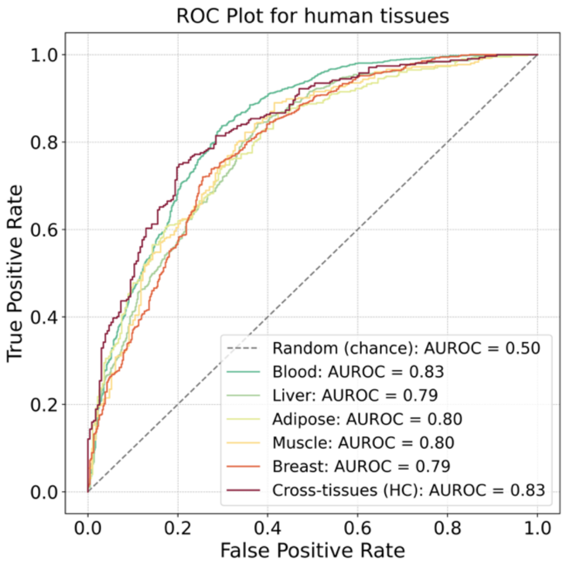


**Supplementary Figure 9** ROC plots and AUROC scores for human models trained and tested with the same number of variants as the corresponding cattle models.

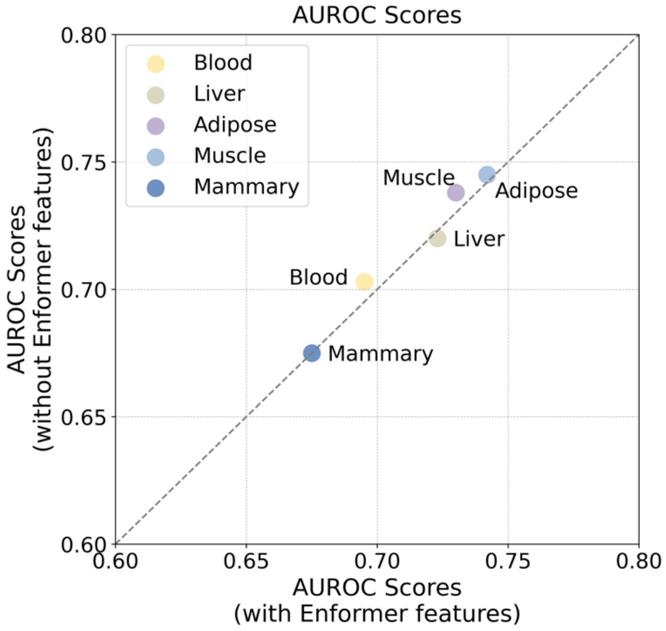


**Supplementary Figure 10** The comparison of the AUROC scores of cattle tissue-specific models trained with or without the full set of Enformer features. The grey dashed line represents parity. The X and Y axis are limited from 0.60 to 0.80 for better clarity of the scatter plot.

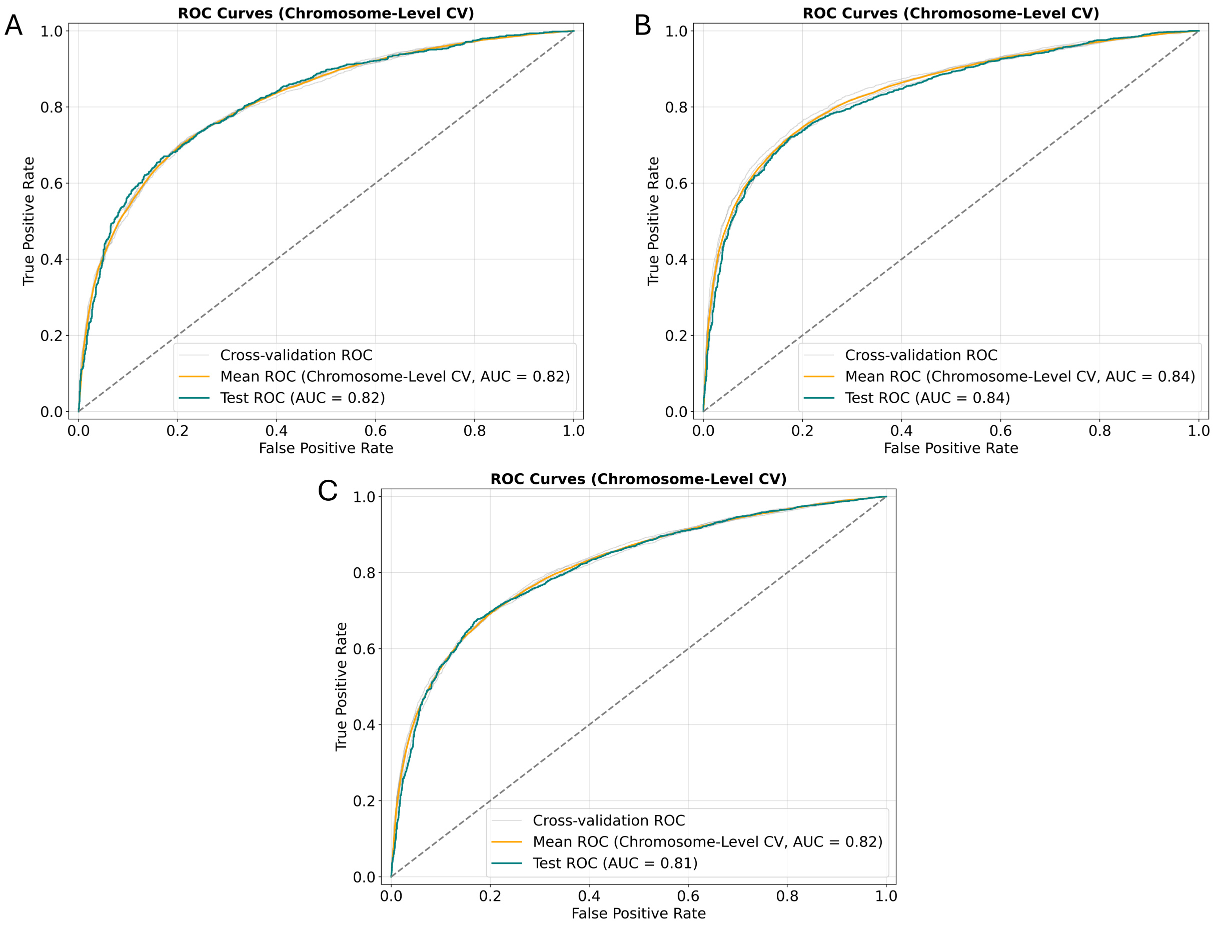

**Supplementary Figure 11** ROC plots and AUROC (AUC) scores for human raQTL models trained and tested in (A) HepG2 cell line. (B) K562 cell line, and (C) both cell lines.

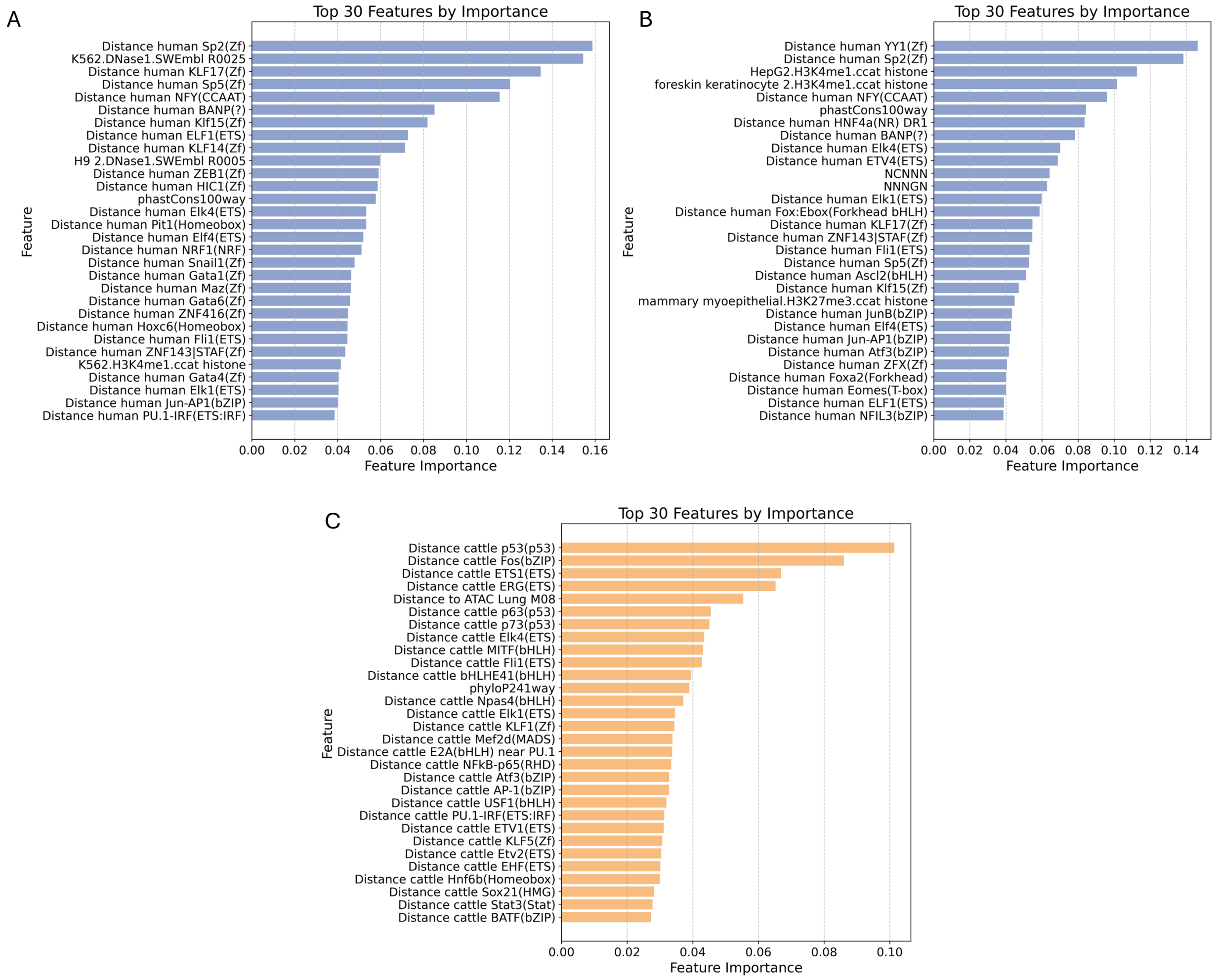


**Supplementary Figure 12 Top 30 features in human and cattle raQTL models (A) K562 cell line, (B) HepG2 cell line and (C) cattle endothelial cell line.** The features are ranked in descending order according to their relative feature importance in the model.

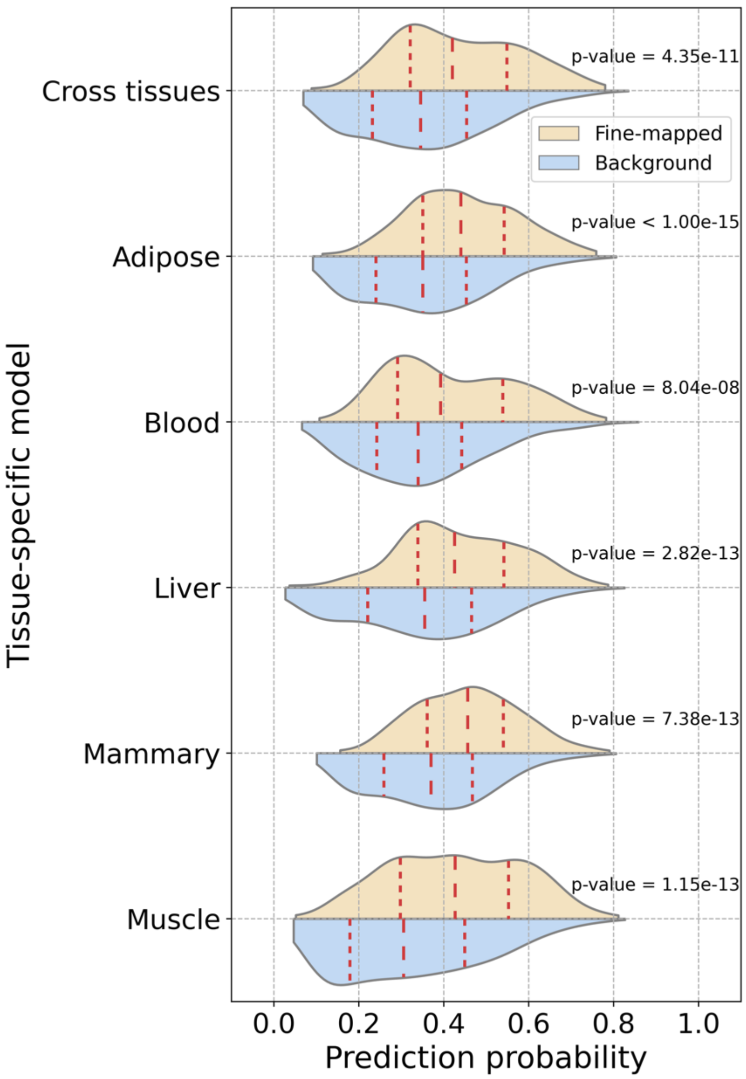


**Supplementary Figure 13 Distributions of model derived prediction probabilities of cattle GWAS fine-mapped and background variants for stature.** The red dashed lines in the violin plots represent the median and quartile values. All prediction probabilities are significantly different between fine-mapped and background groups in each experiment. Their corresponding p-values are displayed in the figure (Two-sample Kolmogorov-Smirnov test).

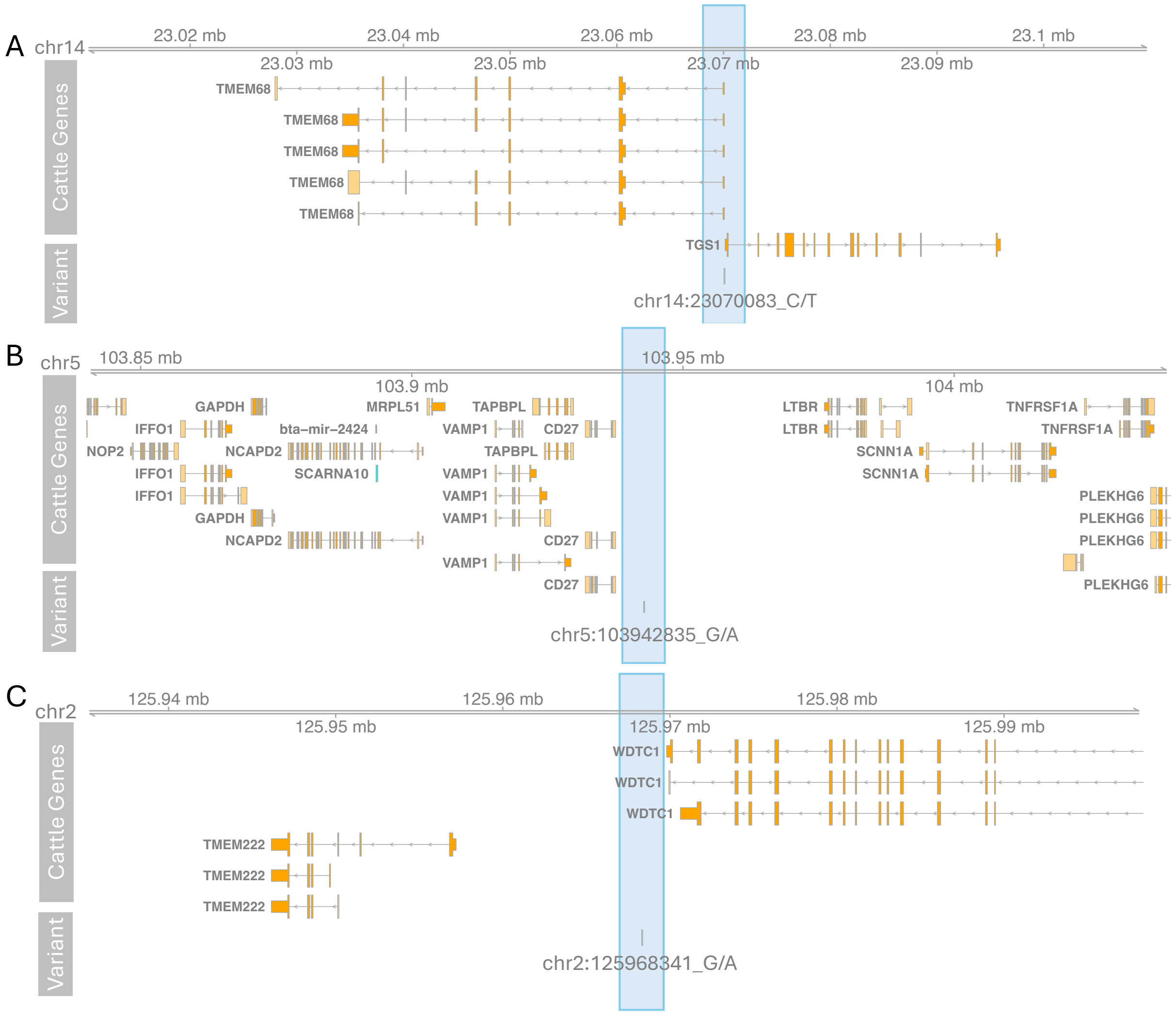


**Supplementary Figure 14 Three cattle trait-associated variants consistently predicted as regulatory variants by all models.** (A) stature-associated variant chr14:23070083_C/T, (B) body weight-associated variant chr5:103942835_G/A, and (C) body condition-associated variant chr2:125968341_G/A.
